# Electron transport chain inhibition increases cellular dependence on purine transport and salvage

**DOI:** 10.1101/2023.05.11.540429

**Authors:** Zheng Wu, Divya Bezwada, Robert C Harris, Chunxiao Pan, Phong T Nguyen, Brandon Faubert, Ling Cai, Feng Cai, Hieu S. Vu, Hongli Chen, Misty Martin- Sandoval, Duyen Do, Wen Gu, Yuannyu Zhang, Bookyung Ko, Bailey Brooks, Sherwin Kelekar, Yuanyuan Zhang, Lauren G Zacharias, K. Celeste Oaxaca, Thomas P Mathews, Javier Garcia-Bermudez, Min Ni, Ralph J. DeBerardinis

**Affiliations:** Children’s Medical Center Research Institute, University of Texas Southwestern Medical Center, Dallas, TX, 75390, USA; Quantitative Biomedical Research Center, Department of Population and Data Sciences, UT Southwestern Medical Center, Dallas, TX, 75390, USA; Simmons Comprehensive Cancer Center, UT Southwestern Medical Center, Dallas, TX, 75390, USA; Department of Medicine, University of Chicago, Chicago, IL, 60637, USA; Department of Radiation Oncology, UT Southwestern Medical Center, Dallas, 75390, TX, USA; Harold C. Simmons Comprehensive Cancer Center, UT Southwestern Medical Center, Dallas, TX, 75390, USA; Howard Hughes Medical Institute, University of Texas Southwestern Medical Center, Dallas, TX, 75390, USA

## Abstract

Cancer cells reprogram their metabolism to support cell growth and proliferation in harsh environments. While many studies have documented the importance of mitochondrial oxidative phosphorylation (OXPHOS) in tumor growth, some cancer cells experience conditions of reduced OXPHOS in vivo and induce alternative metabolic pathways to compensate. To assess how human cells respond to mitochondrial dysfunction, we performed metabolomics in fibroblasts and plasma from patients with inborn errors of mitochondrial metabolism, and in cancer cells subjected to inhibition of the electron transport chain (ETC). All these analyses revealed extensive perturbations in purine-related metabolites; in non-small cell lung cancer (NSCLC) cells, ETC blockade led to purine metabolite accumulation arising from a reduced cytosolic NAD^+^/NADH ratio (NADH reductive stress). Stable isotope tracing demonstrated that ETC deficiency suppressed de novo purine nucleotide synthesis while enhancing purine salvage. Analysis of NSCLC patients infused with [U-^13^C]glucose revealed that tumors with markers of low oxidative mitochondrial metabolism exhibited high expression of the purine salvage enzyme HPRT1 and abundant levels of the HPRT1 product inosine monophosphate (IMP). ETC blockade also induced production of ribose-5’ phosphate (R5P) by the pentose phosphate pathway (PPP) and import of purine nucleobases. Blocking either HPRT1 or nucleoside transporters sensitized cancer cells to ETC inhibition, and overexpressing nucleoside transporters was sufficient to drive growth of NSCLC xenografts. Collectively, this study mechanistically delineates how cells compensate for suppressed purine metabolism in response to ETC blockade, and uncovers a new metabolic vulnerability in tumors experiencing NADH excess.

## INTRODUCTION

Cancer and inborn errors of metabolism (IEMs) are characterized by mutations that perturb cellular metabolism. These diseases provide an opportunity to uncover how metabolic anomalies dysregulate pathways that ultimately lead to tissue dysfunction in humans. Although cancer and IEMs are clinically very different, they share some pathogenic mechanisms. Oncogenic signaling regulates many of the same metabolic pathways that become dysfunctional in IEMs, including glycolysis, amino acid oxidation, the urea cycle and others^1–4^. Some enzymes that are mutated in IEMs, including succinate dehydrogenase and fumarate hydratase, are tumor suppressors in adult-onset cancers^5–7^. Therefore, studying metabolic reprogramming in cancer can provide insights relevant to the pathophysiology of IEMs, and vice versa^8^.

Alterations of mitochondrial function are common in both IEMs and cancer. Mitochondria house critical pathways such as the tricarboxylic acid (TCA) cycle and the electron transport chain (ETC), which are required for oxidative phosphorylation (OXPHOS). These pathways support cell survival and growth by producing energy, biosynthetic precursors, and signaling molecules. Dozens of human IEMs are caused by mutations of mitochondrial enzymes, particularly subunits of the ETC^9^. Tumors have variable mitochondrial activity. Mouse models of cancer indicate that TCA cycle turnover is suppressed in solid tumors at the site of origin relative to adjacent tissue, but that metastases have higher TCA cycle activity^10^. However, eliminating ETC components severely suppresses tumor growth, even at the site of implantation, indicating the requirement for at least a modest amount of ETC activity in those models^11, 12^. Despite the prominence of nonsynonymous point muations in the mitochondrial DNA (mtDNA) of many human cancers, tumors generally do not select for such mutations, and may in fact select against them^13, 14^. Human tumors are variable in the abundance of mtDNA, and a subset of cancer types shows clear evidence for genetic suppression of ETC function^5, 7, 15–17^. Efforts to directly assess tumor metabolism in cancer patients using intra-operative stable isotope infusions have revealed variability in markers of mitochondrial function. Human non-small cell lung cancers (NSCLCs) infused with ^13^C-glucose display variable TCA cycle labeling, implying that there is heterogeneity in how glucose oxidation supplies the TCA cycle in individual tumors^18, 19^. This variation could be related to intrinsic properties of tumor cells^20, 21^, or to environmental factors, such as hypoxia, which may limit oxidation of glucose and other nutrients.

A key marker of ETC dysfunction relevant to cancer cell growth is NADH reductive stress (i.e., low NAD^+^/NADH ratio), which occurs when NADH production outpaces its oxidation to NAD^+^, particularly by ETC Complex I. This form of metabolic stress has broad implications for intermediary metabolism as many NAD(H)-dependent oxidoreductases are sensitive to the NAD^+^/NADH ratio. High demands for NAD^+^ can impose a metabolic bottleneck on growth because some pathways that produce precursors for macromolecular synthesis (e.g. glycolysis, the TCA cycle) require NAD^+^-dependent oxidoreductases^22, 23^. The Warburg effect, which describes the conversion of glucose to lactate in the presence of oxygen observed in most cancer cells, is thought to reflect the need to regenerate NAD^+^ by lactate dehydrogenase when mitochondrial redox shuttles become oversaturated^24^. Identifying consistent metabolic responses when mitochondrial function is compromised and subsequent redox ratios are low may provide new insights about metabolic diseases, including cancer.

In this study, we sought to identify cellular metabolic alterations that occur downstream of mitochondrial dysfunction in humans. Metabolomics revealed purine metabolism as a commonly altered pathway in fibroblasts from patients with a range of defects involving the mitochondria. Surprisingly, purine metabolism was altered as often as the TCA cycle. Purines are produced either through an energetically-demanding pathway of de novo synthesis, or by salvage reactions that regenerate purine nucleotides from bases and nucleosides. Using cancer cell lines, we find that ETC impairments suppress de novo purine synthesis, activate purine salvage, and increase dependence on the salvage enzyme HPRT1, which is dispensable in cells with functional ETCs.

## RESULTS

### Altered purine metabolism is a common feature of human mitochondrial disease

To identify the effects of mitochondrial dysfunction in humans, we first analyzed metabolomics in fibroblasts from patients with IEMs affecting the mitochondria. These patients came from a cohort of individuals being evaluated clinically for IEMs, and their defects involved mitochondrial RNA processing and translation, lipoylation of mitochondrial enzymes, the electron transport chain (ETC) and other processes (Figure 1A). We used metabolite set enrichment analysis (MSEA) to identify pathways perturbed among these fibroblasts compared to fibroblasts from healthy controls. As expected, the TCA cycle was commonly perturbed in fibroblasts from patients with mitochondrial disorders (Figure 1B). Unexpectedly, purine metabolism was even more frequently altered in this cohort (Figure 1C). We also analyzed plasma from a previously-reported girl with deficiency of Lipoyltransferase-1 (LIPT1), the enzyme that adds a lipoyl-AMP cofactor to several enzymes required to maintain flux into and around the TCA cycle, including pyruvate dehydrogenase (PDH) and α-ketoglutarate dehydrogenase (AKGDH)^25^. We compared metabolomics data from this patient’s plasma collected during several hospitalizations to healthy controls. As expected, the LIPT1-deficient patient had elevated lactate, alanine, and α-ketoglutarate, consistent with PDH and AKGDH suppression (Figure 1D). Several purine metabolites were also consistently increased, indicating systemic differences in this pathway from healthy controls (Figure 1E).

**Figure 1.**
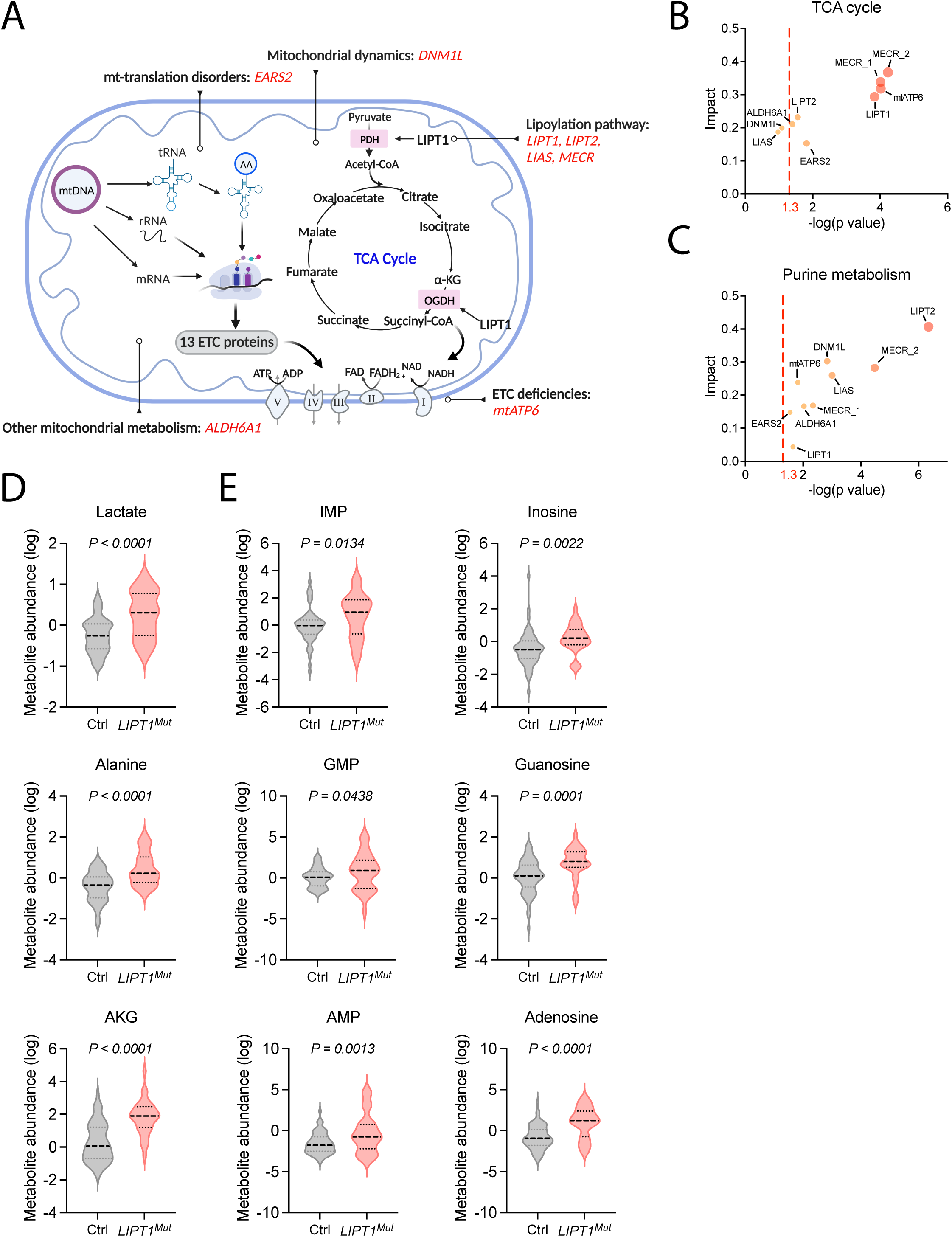
Metabolomic profiling of clinical patients with mitochondrial disorders. **A**. Illustration of the human mitochondrial defects included in the data in panels B-E. **B.** and **C.** Metabolic pathway analysis of the TCA cycle (**B**) and the purine pathway (**C**) in primary fibroblasts from patients with the indicated mitochondrial defects. **D. and E.** Plasma metabolite levels from a patient with LIPT1 deficiency and healthy controls. n = 60 (healthy); n = 28 (LIPT1 deficiency; samples collected on different days). An unpaired, two-sided t test was used for the statistical analysis.

### ETC blockade increases purine metabolites

To assess mitochondrial dysfunction in a simpler system, we treated human H460 NSCLC cells with IACS-010759, a potent mitochondrial ETC complex I inhibitor, to block mitochondrial respiration (Figure 2A)^26^. The drug reduced the cellular NAD^+^/NADH ratio, reflecting blockade of complex I (Figure 2B), and induced a compensatory increase in glucose uptake and lactate secretion (Figures S1A and S1B). Metabolomics revealed widespread effects of IACS-010759, including reduced levels of several metabolites and amino acids related to TCA cycle function (Figure S1C and S1D)^27–29^. Many purine-related metabolites were also altered upon IACS-010759 treatment (Figure 2C). It was notable that several purine nucleotides (e.g. IMP, GMP, and AMP) were elevated despite the depletion of aspartate, a precursor for de novo purine synthesis. These elevations were also observed in other cell lines after treatment with IACS-010759 (Figures S1E and S1F). To test whether IACS-10759 impacts purine metabolism in NSCLC xenografts in vivo, H460 cells were subcutaneously implanted into immunocompromised mice and treated daily with IACS-010759 or vehicle for five days, and samples were processed for metabolomics^30^. Among many alterations, purine metabolism stood out as by far the most affected pathway, with drastic accumulations in GMP and guanosine (Figure S1G-S1I).

**Figure 2.**
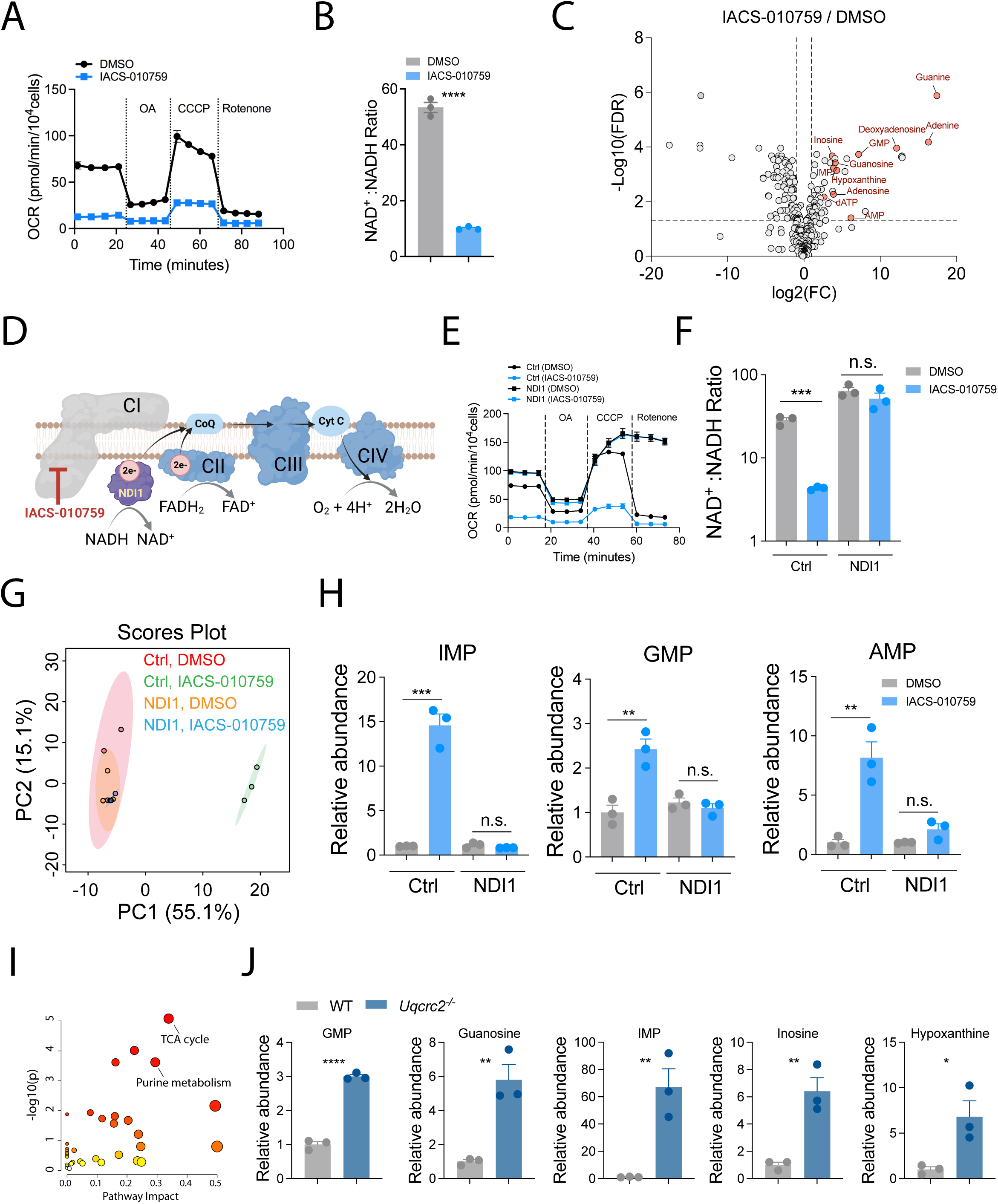
Mitochondrial ETC blockade leads to accumulation of purine metabolites. **A**. Oxygen consumption rates of H460 cells pre-treated with DMSO or 25 nM IACS-010759. OA: oligomycin A. Data represent one of three independent experiments. **B.** NAD^+^/NADH ratio in H460 cells treated with DMSO or 25 nM IACS-010759 for 24 hours. Data are mean from three replicates. **C.** Volcano plot showing metabolomic changes in H460 cells treated with DMSO or 25 nM IACS-010759 for 24 hours. The pink circles are increased purine metabolites with FDR < 0.05. **D.** Schematic illustrating the mechanism of NDI1 in rescue of ETC complex I blockade by IACS-010759. **E.** Oxygen consumption rates of control and NDI1-expressing H460 cells pre-treated with DMSO or 25 nM IACS-010759. OA: oligomycin A. Data represent one of three independent experiments. **F.** NAD^+^/NADH ratio in control and NDI1-expressing H460 cells treated with DMSO or 25 nM IACS-010759 for 24 hours. Data are means from three replicates. **G.** Principal component analysis of metabolomic profiles in control and NDI1-expressing H460 cells treated with DMSO or 25 nM IACS-010759 for 24 hours. **H.** Relative abundance of the indicated purine nucleotides in control and NDI1-expressing H460 cells treated with DMSO or 25 nM IACS-010759 for 24 hours. Data are means from three replicates. **I.** Metabolic pathway analysis of differential metabolites in *UQCRC2*-depleted (*UQCRC2^-/-^*) H460 cells compared to the parental cells. **J.** Relative abundance of the indicated purine metabolites in WT and *UQCRC2^-/-^* H460 cells. Data are means from three replicates. Unpaired, two-sided t tests were used for the statistical analyses. ****: P < 0.0001; ***: P < 0.001, **: P < 0.01, *: P < 0.05; n.s.: P > 0.05. Error bars denote SEM.

To verify that these changes induced by IACS-010759 were caused by complex I inhibition, we expressed *Saccharomyces cerevisiae* alternative NADH dehydrogenase (NDI1), which is resistant to IACS-010759 (Figure 2D). NDI1 boosted basal mitochondrial respiration in H460 cells and rendered them resistant to IASC-010759 and rotenone (Figure 2E). NDI1 expression also normalized the NAD^+^/NADH ratio, glucose uptake and lactate secretion in the presence of IACS-010759 (Figures 2F, S2A). Control H460 cells expressing an empty vector displayed growth suppression upon IACS-010759 treatment, and extensive cell death in medium containing galactose instead of glucose. These effects were reversed by NDI1 (Figure S2B). Metabolomics revealed that nearly all IACS-010759-induced metabolic alterations, including those involving purines, were corrected by NDI1 expression (Figures 2G and 2H, S2C and S2D). Of note, MSEA identified purine metabolism as the top-scoring pathway from metabolites that were altered by IACS-010759 in control but not NDI1-expressing cells (Figure S2E), identifying this pathway as the most responsive to complex I blockade.

Since the ETC involves multiple complexes in addition to complex I, we also generated an isogenic H460 cell line depleted for UQCRC2, a component of ETC complex III (*UQCRC2^-/-^* cells, Figure S3A). UQCRC2 ablation impaired mitochondrial respiration (Figure S3B), suppressed cell growth (Figure S3C), and induced a distinct metabolomic profile (Figures S3D and S3E), including marked purine accumulation (Figures 2I and 2J).

### Cytosolic NAD^+^/NADH ratio regulates purine metabolism upon ETC blockade

Mitochondrial respiration is coupled to oxidation of reducing equivalents (i.e., NADH and FADH_2_), ATP synthesis, and mitochondrial membrane potential, all of which are suppressed by IACS-010759 and rescued by NDI1. It was unclear which aspect of mitochondrial function modulates purine abundance. Given that NAD^+^ is critical for various oxidoreductase reactions, we reasoned that the decreased NAD^+^/NADH ratio impacts purine metabolism when mitochondrial ETC is compromised. In order to dissociate NADH oxidation from OXPHOS, we expressed the water-forming NADH oxidase from *Lactobacillus brevis* (*Lb*NOX) that utilizes oxygen to convert NADH to NAD^+^ (Figure 3A)^31^. We localized *Lb*NOX to cytosol or mitochondria in *UQCRC2^-/-^*cells to ameliorate NADH accumulation in either compartment (Figures 3B and 3C)^31^. Mito-*Lb*NOX modestly increased the total cellular NAD^+^/NADH ratio, whereas Cyto-*Lb*NOX did not have an appreciable effect in whole cell lysates (Figure 3D). Nevertheless, consistent with previous studies^31^, both versions of *Lb*NOX improved *UQCRC2^-/-^* cell growth in the absence of pyruvate and uridine, and the effect was more pronounced with Cyto-*Lb*NOX (Figure S3F). Importantly, in *UQCRC2^-/-^* cells, *Lb*NOX did not significantly affect oxygen consumption, indicating that LbNOX increases the NAD^+^/NADH ratio independently of OXPHOS (Figure 3E). We assessed metabolomic profiles of WT and *UQCRC2^-/-^* cells expressing either an empty vector (EV) or Mito/Cyto-*Lb*NOX, all grown in medium lacking pyruvate and uridine supplementation. Under these conditions, Cyto-*Lb*NOX had a larger impact on the metabolic profile than Mito-*Lb*NOX (Figure 3F). Interestingly, Cyto-*Lb*NOX but not Mito-*Lb*NOX reduced IMP and hypoxanthine levels in *UQCRC2^-/-^* cells (Figure 3G), and this correlated with increased restoration of aspartate by Cyto-*Lb*NOX (Figure S3G), although neither Cyto-*Lb*NOX nor Mito-*Lb*NOX completely resolved the metabolic effects of *UQCRC2^-/-^* loss (Figure S3H). Thus, changes in purine pathway likely stem from cytosolic redox imbalance even when the inciting event is ETC blockade. To further test this hypothesis, we supplemented the medium with α-ketobutyrate (AKB), which has been used to alleviate cytosolic NADH accumulation (Figure 3H)^28^. Similar to Cyto-*Lb*NOX expression, AKB treatment reduced hypoxanthine and IMP levels and enhanced cell growth (Figures 3I and 3J). These data corroborate a previous finding showing that pyruvate supplementation does not correct the mitochondrial NAD^+^/NADH ratio in ETC-deficient cells, but does restore aspartate abundance and cell growth^32^.

**Figure 3.**
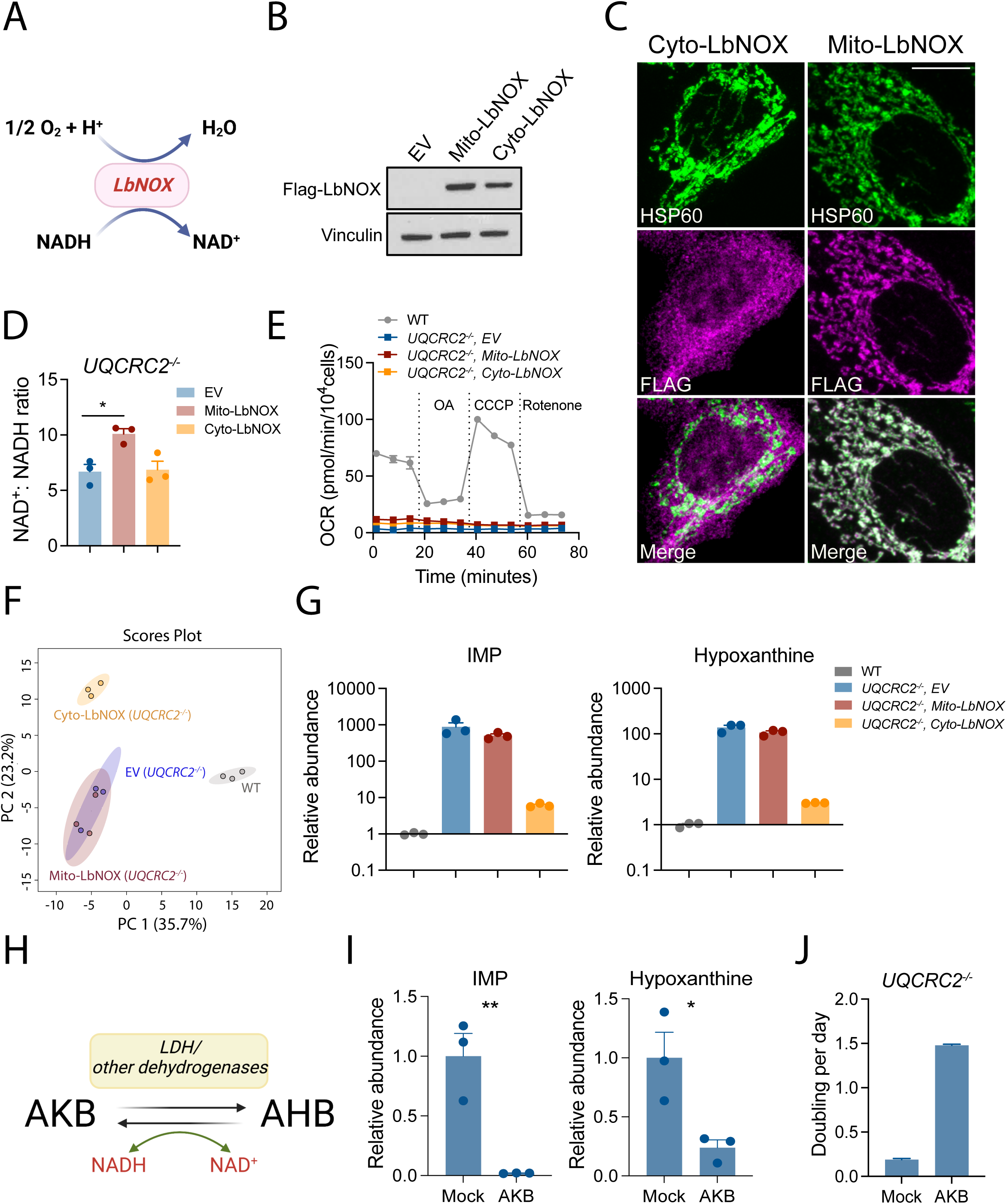
Cytosolic NAD(H) imbalance contributes to purine accumulation in ETC deficient cells. **A**. Schematic of *Lb*NOX-catalyzed reaction. **B.** Western blot validating expression of Flag-tagged *Lb*NOX in *UQCRC2^-/-^* H460 cells. Vinculin is the loading control. **C.** Immunofluorescence showing the subcellular localizations of the indicated Flag-tagged LbNOX proteins. HSP60 serves as a mitochondrial matrix marker. Scale bar represents 10 µm. **D.** NAD^+^/NADH ratio in *UQCRC2^-/-^* cells expressing an empty vector (EV), Mito-*Lb*NOX or Cyto-*Lb*NOX. Data are means from three replicates. **E.** Oxygen consumption rates of WT and *UQCRC2^-/-^* cells expressing EV, Mito*-Lb*NOX or Cyto*-Lb*NOX. OA: oligomycin A. Data are from one of three independent experiments. **F.** Principal component analysis of metabolomic profiles in WT and *UQCRC2^-/-^* cells expressing EV, Mito*-Lb*NOX, or Cyto*-Lb*NOX. **G.** Comparison of the indicated purine metabolites in WT and *UQCRC2^-/-^* cells expressing EV, Mito*-Lb*NOX, or Cyto*-Lb*NOX. Data are means from three replicates. **H.** Schematic illustrating how AKB mitigates cytosolic NADH reductive stress. **I.** Relative abundance of IMP and hypoxanthine in *UQCRC2^-/-^* cells cultured with or without 1 mM AKB. Data are mean from three replicates. **J.** Growth rates of *UQCRC2^-/-^*cells cultured with or without 1 mM AKB (n=12). Data are from one of three independent experiments. Unpaired, two-sided t tests were used for the statistical analyses. **: P < 0.01, *: P < 0.05. Error bars denote SEM.

### ETC blockade suppresses de novo purine synthesis

We next defined how suppressing respiration altered purine levels. IACS-010759 nearly eliminated 5-Aminoimidazole-4-carboxamide ribonucleotide (AICAR) and Phosphoribosylaminoimidazolesuccinocarboxamide (SAICAR), two intermediates in the de novo purine synthesis pathway (Figure 4A). To more rigorously assess de novo purine synthesis, we conducted kinetic [amide-^15^N]glutamine tracing in H460 cells with or without IACS-010759. The de novo pathway incorporates two and three ^15^N nuclei into IMP and GMP, respectively (Figure 4B). Compared to untreated cells, control cells treated with IACS-010759 exhibited lower fractional enrichment of m+2 IMP and m+3 GMP throughout the time course, and this effect was reversed by NDI1 (Figure 4C). IACS-010759 also suppressed both m+2 labeling and abundance of AICAR in control but not NDI1-expressing cells (Figure 4D). As expected if the changes in purine metabolism arose as a direct consequence of ETC blockade, IACS-010759 had little effect on expression of enzymes in the de novo pathway (Figure S4A).

**Figure 4.**
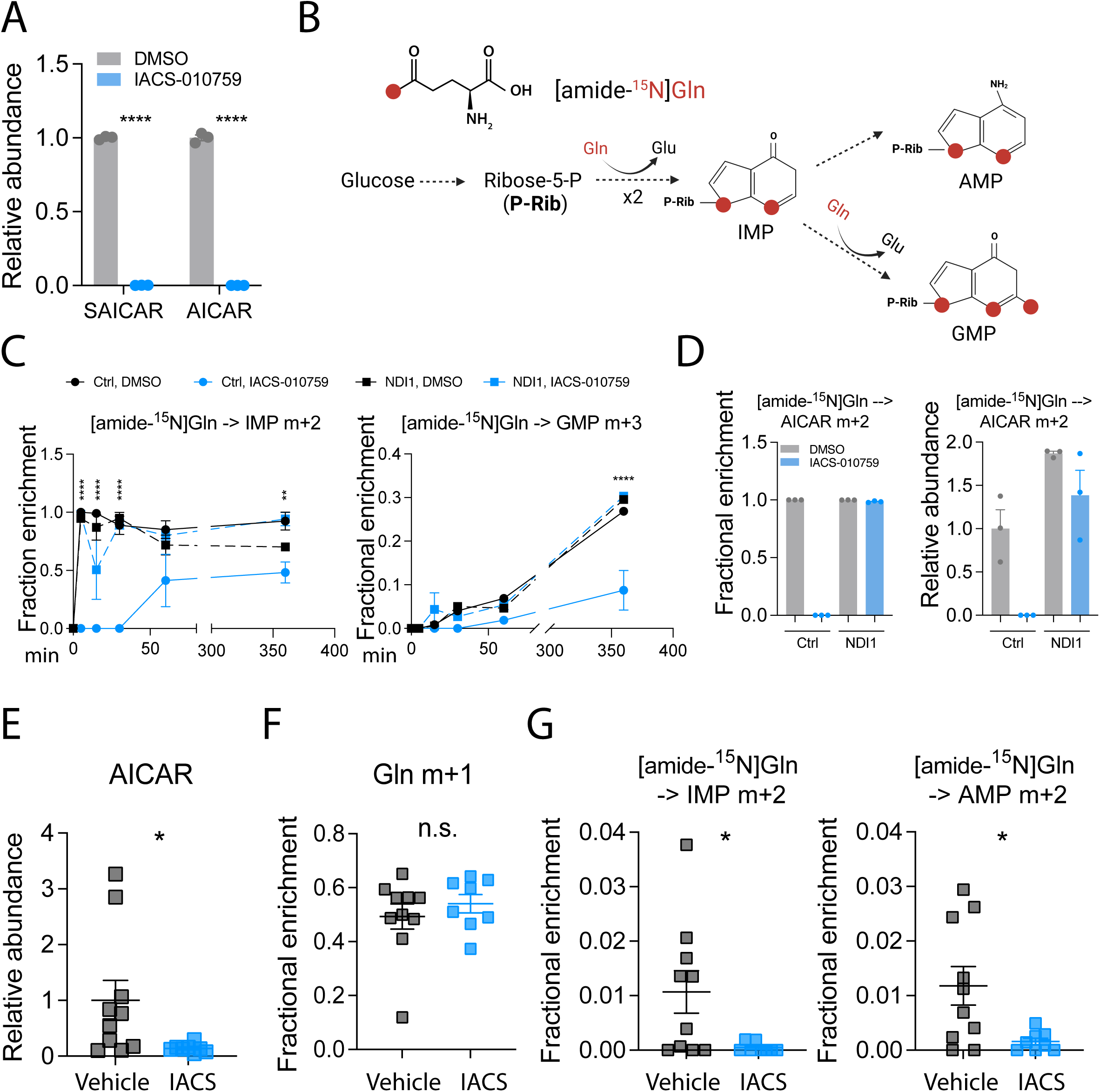
ETC blockade suppresses de novo purine nucleotide synthesis. **A**. Relative abundance of SAICAR and AICAR in H460 cells treated with DMSO or 25 nM IACS-010759 for 24 hours. Data are means from three replicates. **B.** Schematic illustrating ^15^N labeling from [amide-^15^N]glutamine during de novo purine nucleotide synthesis. **C.** Time-dependent fractional enrichment of m+2 IMP and m+3 GMP in control and NDI1-expressing H460 cells pre-treated with DMSO or 25 nM IACS-010759 for 24 hours. Data are means from three replicates at each time point. **D.** Fractional enrichment and relative abundance of m+2 AICAR after 5 minutes of culture with [U-^13^C]glucose in control and NDI1-expressing H460 cells pre-treated with DMSO or 25 nM IACS-010759 for 24 hours. Data are means from three replicates. **E.** Relative abundance of AICAR in H460 xenografts treated with vehicle or IACS-010759 for 5 days. Vehicle (n=10), IACS (n=8). **F-G.** Fractional enrichment of m+1 glutamine (**F**) m+2 IMP and m+2 AMP (**G**) in vehicle and IACS-010759-treated H460 xenografts after 4 hours of [amide-^15^N]glutamine infusion. Vehicle (n=10), IACS (n=8). Unpaired, two-sided t tests (**A**, **D**, and **E-G**) and two-way ANOVA (**C**) were used for the statistical analyses. ****: P < 0.0001; **: P < 0.01, *: P < 0.05; n.s.: P > 0.05. Error bars denote SEM.

To test whether ETC blockade impacts purine metabolism in tumors in vivo, H460 xenograft-bearing mice were dosed with IACS-010759 daily for five days and then infused via tail vein with [amide-^15^N]glutamine for four hours. AICAR abundance declined in tumors treated with IACS-010759 (Figure 4E). IACS-010759 had no effect on tumor enrichment of m+1 glutamine (Figure 4F) but it suppressed m+2 labeling in IMP and AMP (Figure 4G), indicating suppressed de novo purine synthesis in vivo.

### Mitochondrial ETC deficiency enhances purine salvage

These results raised the question of how cells manage to elevate purine nucleotides under conditions of ETC blockade and a low NAD^+^/NADH ratio. To address this question, we cultured cells with uniformly ^13^C-labeled glucose (i.e., [U-^13^C]glucose) to examine glucose’s contributions to purines in the presence and absence of IACS-010759. While de novo purine synthesis yields various purine nucleotide isotopologues reflecting labeling in both the ribose backbone and the purine base, purines produced from the salvage pathway are dominated by m+5 labeling from ribose without any additional labeling from the base (Figure 5A). Vehicle-treated cells displayed the expected heterogeneity in IMP isotopologues, but labeling was almost entirely m+5 in IACS-010759-treated control cells. Cells expressing NDI1 resisted this effect of IACS-010759 (Figure 5B). Similar effects occurred in GTP and ATP, although overall labeling was lower than IMP (Figure S4B). These labeling patterns indicate that complex I blockade increases the contribution of the purine salvage pathway to purine pools. IACS-010759 increased time-dependent m+5 labeling of both IMP and GMP (Figure 5C).

**Figure 5.**
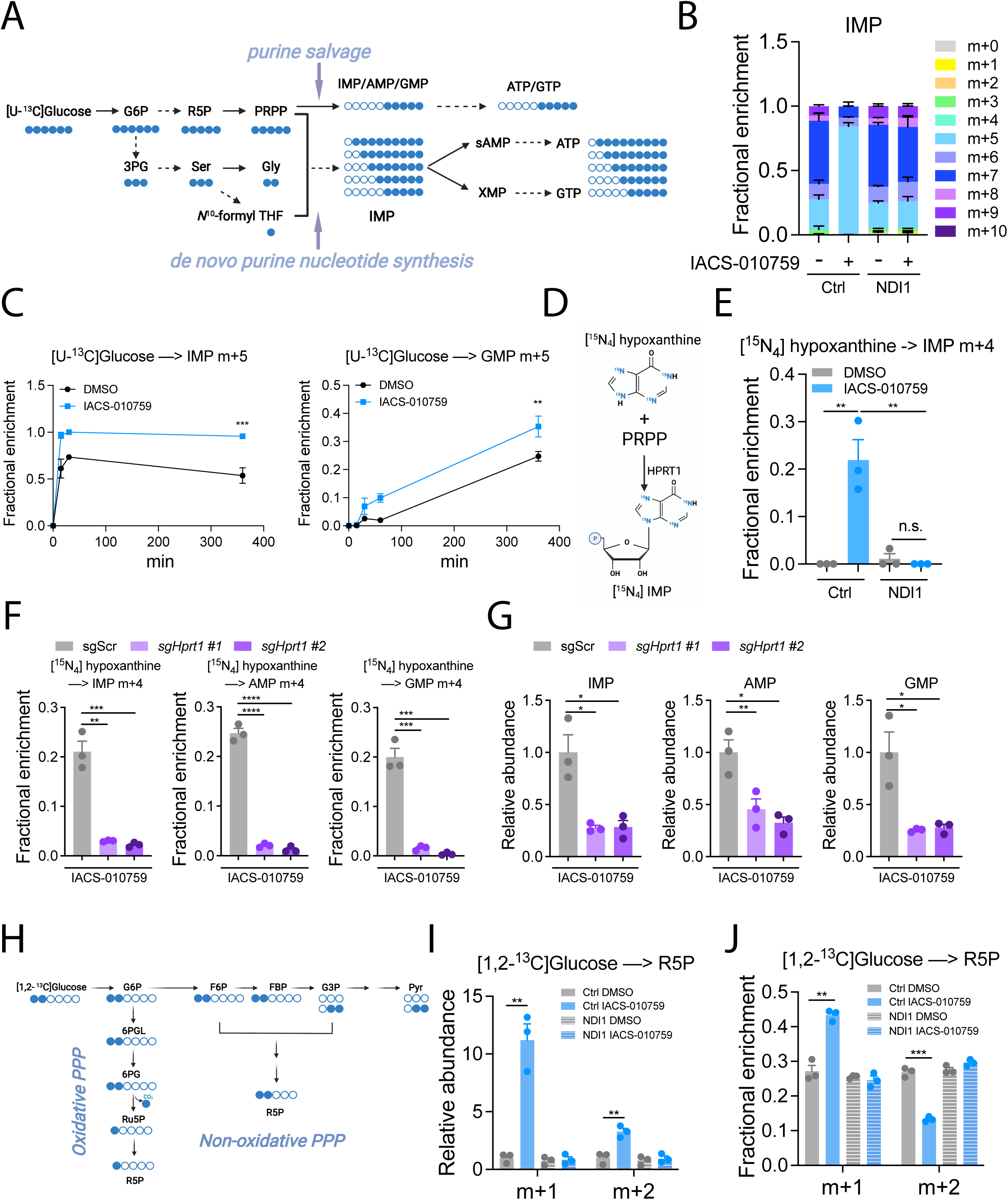
ETC blockade promotes purine salvage. **A**. Schematic illustrating ^13^C labeling of purines from [U-^13^C]glucose. **B.** ^13^C labeling in IMP after 6 hours of culture with [U-^13^C]glucose in control and NDI1-expressing H460 cells pre-treated with DMSO or 25 nM IACS-010759 for 24 hours. Data are means from three replicates. **C.** Time-dependent labeling of m+5 IMP and m+5 GMP in H460 cells pre-treated with DMSO or 25 nM IACS-010759 for 24 hours prior to culture with [U-^13^C]glucose. Data are means from three replicates. **D.** Schematic illustrating conversion of [^15^N_4_]hypoxanthine to m+4 IMP during HPRT1-mediated salvage. **E.** Fractional enrichment of m+4 IMP from [^15^N_4_]hypoxanthine during 6 hours of tracing in control and NDI1-expressing H460 cells pre-treated with DMSO or 25 nM IACS-010759 for 24 hours. **F.** Fractional enrichment of m+4 IMP, AMP, and GMP in control (*sgScr*) or HPRT1-depleted (*sgHPRT1*) cells after 24 hours of pre-treatment with 25 nM IACS-010759 followed by 6 hours of culture with [^15^N_4_]hypoxanthine. Data are means from three replicates. **G.** Relative abundance of purine nucleotides in control (sgScr) and HPRT1-depleted H460 cells with 24 hours of IACS-010759 treatment. Data are means from three replicates. **H.** Schematic illustrating ^13^C labeling of R5P from [1,2-^13^C]glucose. **I-J.** Relative abundance (**I**) and fractional enrichment (**J**) of m+1 and m+2 R5P after 6 hours of culture in [1,2-^13^C]glucose. Control and NDI1-expressing H460 cells were pre-treated with DMSO or IACS-010759 for 24 hours. Data are means from three replicates. Unpaired, two-sided t tests (**E**-**G**), multiple t test (**I** and **J**), and two-way ANOVA (**C**) were used for the statistical analyses. ****: P < 0.0001; ***: P < 0.001; **: P < 0.01, n.s.: P > 0.05. Error bars denote SEM.

To examine the purine salvage pathway directly, we traced nitrogen-labeled hypoxanthine (i.e., [^15^N_4_]hypoxanthine) in cells treated with vehicle or IACS-010759 (Figure 5D). After 6 hours of culture with [^15^N_4_]hypoxanthine, IACS-010759-treated control cells had higher m+4 IMP enrichment than untreated cells, with NDI1 again reversing this effect (Figure 5E). Kinetic tracing revealed increased labeling of both AMP and GMP from [^15^N_4_]hypoxanthine in IACS-010759-treated cells (Figure S4C). To test whether purine salvage was necessary for purine nucleotide accumulation during complex I blockade, we utilized CRISPR-Cas9 with two independent guide RNAs (*sgHprt1 #1* and *#2*) to generate H460 cells defective in the purine salvage enzyme hypoxanthine-guanine phosphoribosyltransferase (HPRT1, Fig. S4D). During [^15^N_4_]hypoxanthine tracing, ablation of HPRT1 led to an elevation in m+4 labeling in intracellular hypoxanthine, but nearly eliminated enrichment of m+4 purine nucleotides induced by IACS-010759 (Figure S4E and 5F). The accumulation of IMP, AMP, and GMP induced by IACS-010759 was also blunted in the absence of HPRT1 (Figure 5G), indicating that HPRT1-mediated purine salvage accounts, at least partially, for the accumulated purine monophosphates upon ETC blockade. Neither HPRT1 enzymatic activity nor its expression was changed in response to defective mitochondrial respiration (Figure S4A and S4F).

### ETC blockade enhances the PPP

Purine salvage requires purine nucleobases and phosphoribosyl diphosphate (PRPP), an activated form of ribose-5-phosphate (R5P) produced in the pentose phosphate pathway (PPP). Therefore, we examined the effects of complex I inhibition on the PPP. After 24 hours of treatment of IACS-010759, R5P pools were relatively depleted in IACS-010759-treated cells (Figure S4G). However, after addition of fresh medium containing [U-^13^C]glucose, both the abundance and m+5 labeling of R5P rose faster in IACS-010759-treated than DMSO-treated cells, indicating rapid synthesis of R5P from glucose during complex I blockade (Figures S4H to S4J). IACS-010759-treated cells also displayed higher enrichment of m+6 6-phosphogluconate (6-PG) and m+7 sedoheptulose 7-phosphate (S7P), two PPP intermediates (Figure S4K). To assess whether the oxidative or non-oxidative branch of the PPP predominated in these cells, we used [1,2-^13^C]glucose as a tracer. In this tracing scheme, both m+1 and m+2 R5P are produced, with m+1 arising predominantly from the oxidative branch and m+2 arising from the non-oxidative branch (Figure 5H). IACS-010759 treatment increased both m+1 and m+2 R5P abundance, but the fractional enrichment of m+1 increased while m+2 decreased in response to IACS-010759 (Figures 5I and 5J). IACS-010759 had no effect on R5P labeling or abundance in NDI1-expressing cells (Figures 5I and 5J). Taken together, the data indicate an activation of PPP activity, primarily through the oxidative branch, in response to ETC blockade.

### HPRT1 is important for NSCLC growth especially when ETC is impaired

Since defective mitochondrial respiration inhibits de novo purine nucleotide synthesis, we hypothesized that blockade of purine salvage is detrimental to ETC-deficient cells. Indeed, while HPRT1 is dispensable for proliferation of ETC-competent cells, cells lacking HPRT1 could not proliferate when treated with IACS-010759 (Figure 6A). To assess HPRT1’s role in purine metabolism in vivo, we subcutaneously injected control and HPRT1-deficient H460 cells into immunocompromised mice and dosed the tumor-bearing mice with vehicle or IACS-010759. HPRT1 deficiency reduced tumor growth even without IACS-010759 treatment, but HPRT1-deficient tumors were more sensitive to IACS-010759 treatment (Figure 6B). These data indicate that although H460 cells tolerate HPRT1 loss in culture, this enzyme is required for maximal tumor growth in vivo and complex I blockade increases sensitivity to HPRT1 loss both in culture and in vivo.

**Figure 6.**
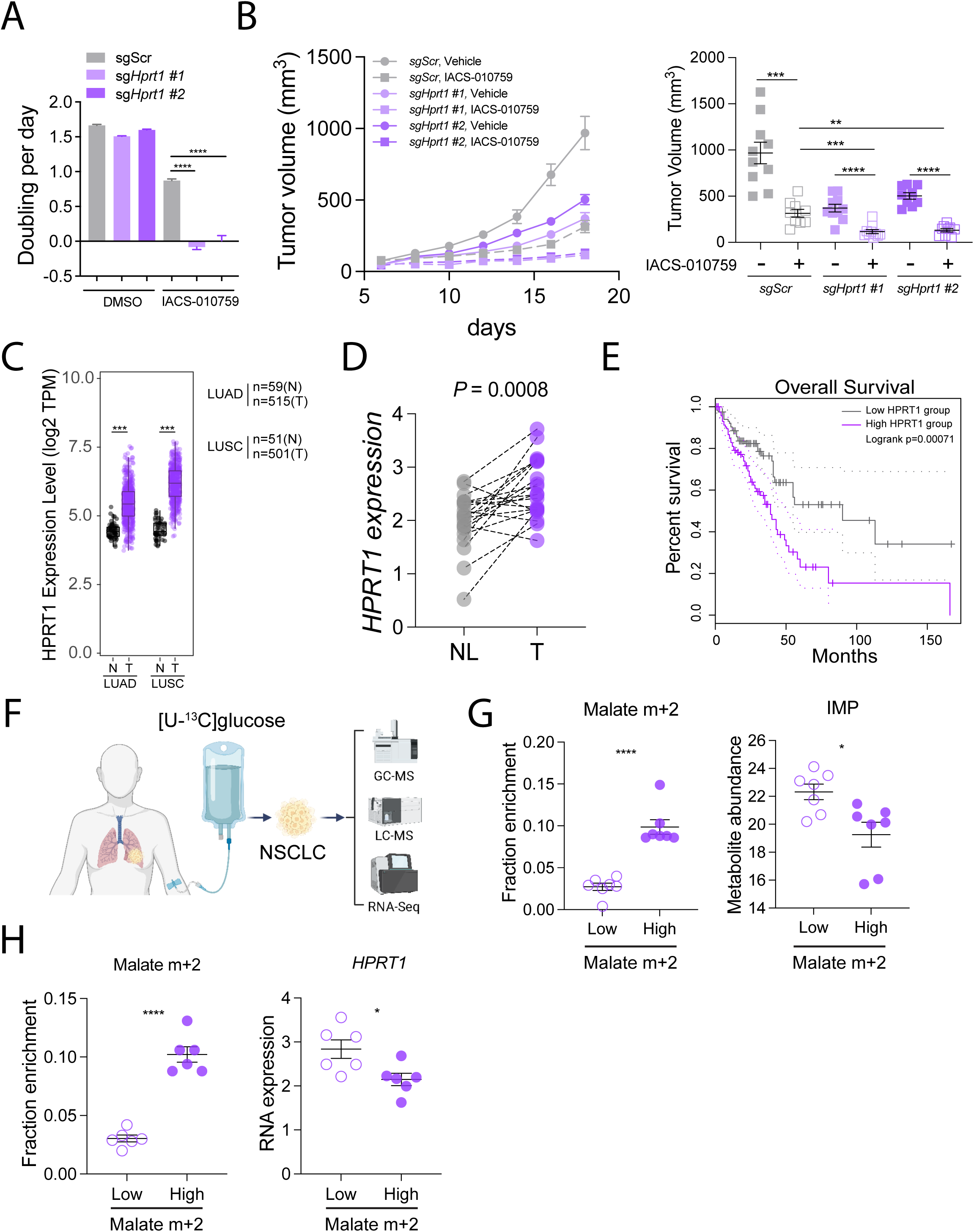
HPRT1 supports NSCLC growth during ETC inhibition. **A**. Cell growth rates of control and HPRT1-depleted H460 cells treated with DMSO or 25 nM IACS-010759 (n=6). Data are from one of three independent experiments. **B.** Subcutaneous tumor growth of control and HPRT1-deficient H460 xenografts treated with vehicle or 5 mg/kg IACS-010759. The right panel shows individual tumor sizes on the day when the tumors were harvested. n=10 for sgScr Vehicle, sgHprt1 #1 Vehicle, and sgHprt1 #1 IACS-010759. n=9 for sgScr IACS-010759, sgHprt1 #2 Vehicle, and sgHprt1 #2 IACS-010759. **C.** *HPRT1* mRNA levels in human LUAD and LUSC tumors (T) or nonmalignant lung tissue (N). Data and statistics were generated using TIMER2.0^68, 69^. **D.** Patient-matched *HPRT1* expression in human NSCLC tumors (T) and adjacent, nonmalignant lung (NL) (n=20). **E.** Kaplan-Meier plot showing overall survival of LUAD patients with high (top 25%, n=120) and low (bottom 25%, n=120) *HPRT1* expression. Hazard ratio (high) = 2.1. p(HR)=0.00097. The plot and statistics were generated using GEPIA 2^70^. **F.** Schematic illustrating intra-operative [U-^13^C]glucose infusion in NSCLC patients followed by tumor resection and multi-omics analyses. **G.** Fractional enrichment of m+2 malate and relative IMP abundance in tumors displaying low or high malate labeling. The analysis was performed on the top and bottom 25% of tumors for malate m+2 labeling (n=7 tumors each with both isotope tracing and metabolomics analysis). **H.** Fractional enrichment of m+2 malate and *HPRT1* mRNA levels in tumors displaying low or high malate labeling. The analysis was performed on the top and bottom 25% of tumors for malate m+2 labeling (n=6 tumors each with both isotope tracing and RNA-Seq analysis). Unpaired, two-sided t tests (**B, G,** and **H**), and a paired t test (**D**) were used for the statistical analyses. ****: P < 0.0001; ***: P < 0.001; **: P < 0.01, n.s.: P > 0.05. Error bars denote SEM.

While de novo purine synthesis is the target of multiple chemotherapeutic drugs, the role of purine salvage in tumor growth remains underappreciated. Analysis of the cancer genome atlas (TCGA) showed higher expression of *HPRT1* in human lung adenocarcinomas and squamous cell carcinomas compared to nonmalignant lung (Figure 6C). We also observed enhanced *HPRT1* expression in NSCLCs relative to patient-matched lung tissue from our own cohort (Figure 6D). High expression of *HPRT1* correlates with poor overall survival of NSCLC patients (Figure 6E).

We next examined how mitochondrial function affects purine metabolism in human NSCLC in vivo. Intra-operative infusion of [U-^13^C]glucose during surgical NSCLC resection leads to variable labeling in TCA cycle intermediates extracted from the tumors (Figure 6F)^18, 19^. In xenografts, ETC activity within NSCLC cells contributes to TCA cycle intermediate labeling^30^, so for this analysis we asked whether labeling of these intermediates is related to markers of purine metabolism. In human NSCLCs, there was a strong correlation between m+2 glutamate and m+2 malate, indicating label propagation around the TCA cycle (Figure S5A). Analysis of ^13^C labeling and metabolite abundance revealed that tumors with low malate m+2 enrichment (bottom quartile, n=7) contained high IMP abundance (Figure 6G). RNA-sequencing data (n=6 tumors with both RNA-sequencing and ^13^C analyses) revealed that tumors with low malate m+2 enrichment exhibited high *HPRT1* expression (Figure 6H). Similar results were also obtained if we used m+2 glutamate for the analyses (Figure S5B and S5C). These data may indicate an enhanced propensity for purine salvage when glucose-dependent labeling of TCA cycle intermediates is low, as would be the case if OXPHOS is relatively impaired.

### Purine nucleotide accumulation induced by complex I inhibition is independent of macroautophagy

We next explored how cells acquire purine nucleobases for the salvage reaction. Cancer cells can exploit autophagy to generate purine nucleotides^33^. Some 80% of cellular RNA is ribosomal RNA, which accounts for most of the ribosomal mass^34, 35^. The selective degradation of ribosomes by autophagy (ribophagy) contributes to nucleotide pools during nutrient starvation, and this process is negatively regulated by mTORC1^36, 37^. Consistent with a previous study^29^, we observed repressed mTORC1 signaling upon IACS-010759 treatment, and this was restored by NDI1 (Figures S6A and S6B). Since mTORC1 inhibits ribophagy, we tested whether mTORC1 suppression induces ribosomal degradation and supplies nucleobases for purine salvage during IACS-010759 treatment. We generated an H460 ribophagy reporter cell line that expresses ribosomal protein 3 (RPS3) fused with a Keima-Red protein (Figure S6C). During ribophagy, RPS3-Keima-Red is cleaved to release Keima protein (Figure S6D, lower band)^36, 38^. Treatment with the mTORC1 inhibitor Torin1 led to the expected cleavage of RPS3-Keima, and this was prevented by the autophagy inhibitor Bafilomycin A (Figure S6D). In contrast, IACS-010759 did not induce ribophagy in this assay (Figure S6D). IACS-010759 decreased p62/SQSTM1, which is indicative of augmented macroautophagy flux (Figure S6D). To examine whether macroautophagy contributes to purine accumulation, we utilized CRISPR-Cas9 technology to delete ATG5 and ATG7, two essential autophagy factors (Figure S6E). However, IACS-010759 resulted in similar IMP, GMP, and AMP levels in control, ATG5-deficient, and ATG7-deficient cells (Figure S6F). Therefore, we conclude that blocking complex I results in purine nucleotide accumulation independently of macroautophagy or ribophagy.

### ETC blockade stimulates purine nucleobase uptake to provide purine nucleotides for cell growth

Metabolic stress induces nutrient scavenging from the microenvironment to sustain cell survival and growth. ETC-deficient cells rely on environmental lipids for cell growth, and pancreatic cancer cells use micropinocytosis when aspartate synthesis is disrupted^39, 40^. Of note, an unbiased CRISPR screen in pancreatic cancer cells identified both HPRT1 and SLC29A1, a nucleoside/base transporter, as conditionally essential during ETC blockade^40^. Consistent with this result, supplementing the culture medium with purine nucleosides (e.g. inosine, guanosine, and adenosine) that can be converted to nucleobases to feed the salvage pathway improved cell growth in the presence of IACS-010759 (Figure S7A). This led us to hypothesize that ETC-deficient cells take up nucleosides or nucleobases from the medium to supply the salvage pathway.

There are two main nucleoside transporter groups, the SLC28 and SLC29 families. Both families can transport most purine nucleosides and nuleobases^41^. From RNA-seq data, we determined that H460 cells only express appreciable levels of SLC29A1 and SLC29A2 (Figure S7B). Treating cells with IACS-010759 induced hypoxanthine uptake, which was blunted by concomitant incubation with the SLC29A1/SLC29A2 inhibitor nitrobenzylthioinosine (NBMPR, Figure S7C). NBMPR reduced labeling of cellular purine nucleotides cultured with [^15^N_4_]hypoxanthine (Figure 7A), and nearly eliminated purine nucleotide accumulation induced by IACS-010759 (Figure 7B).

**Figure 7.**
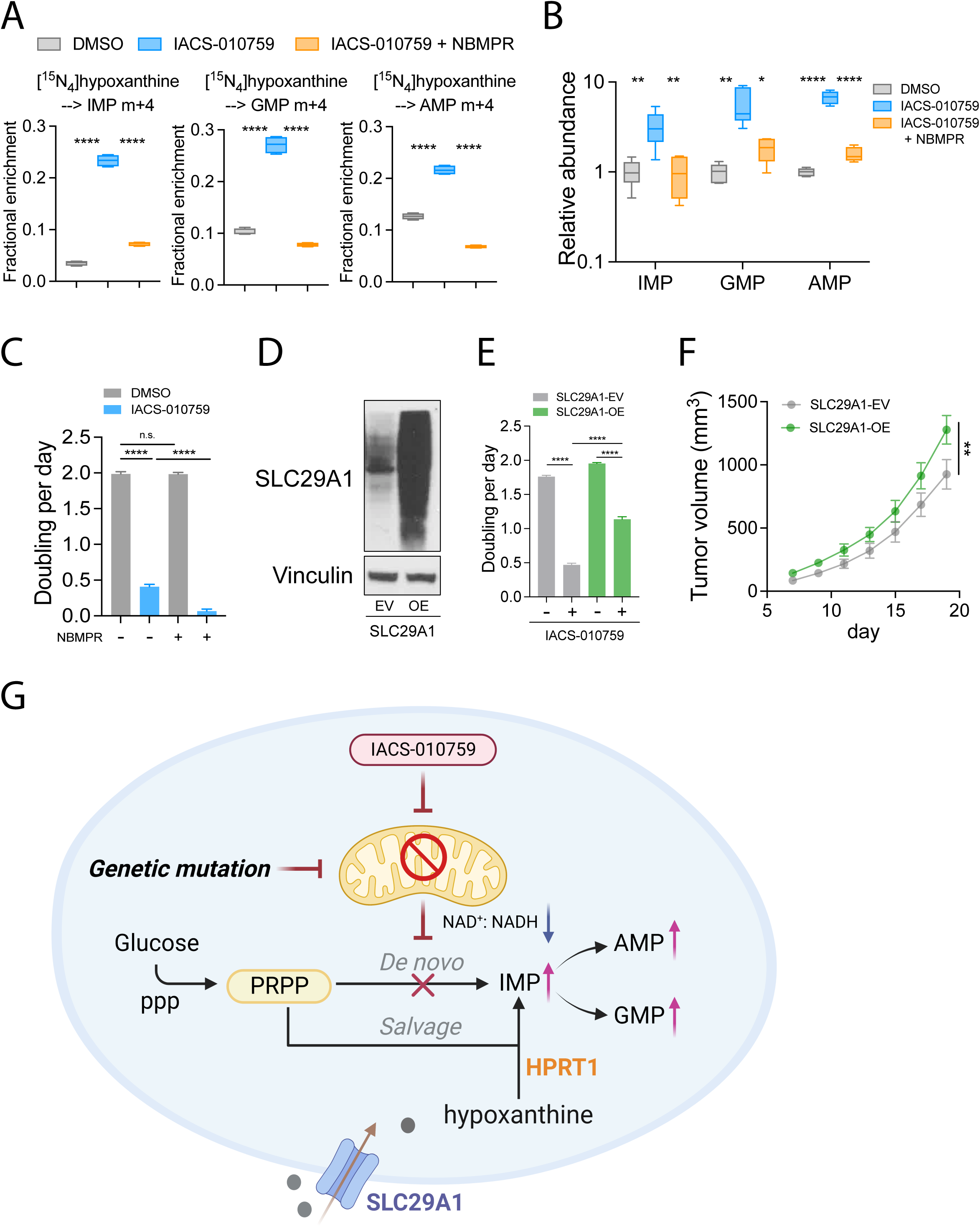
Purine uptake is required to supply salvage upon ETC blockade. **A**. Fractional enrichment of m+4 IMP, GMP, and AMP after 6 hours of culture with [^15^N_4_]hypoxanthine, following 24-hours of treatment with DMSO or 25 nM IACS-010759, with or without 50 µM NBMPR. Data are means from four replicates. **B.** Relative abundance of the indicated purine nucleotides in H460 cells after 24 hours of treatment with DMSO or 25 nM IACS-010759, with or without 50 µM NBMPR. Data are means from six replicates. **C.** Growth rates of H460 cells treated with 50 µM NBMPR, 25 nM IACS-010759, or both (n=8). Data are from one of three independent experiments. **D.** Western blot validating overexpression of SLC29A1. Vinculin is the loading control. **E.** Growth rates of control (SLC29A1-EV) and SLC29A1-overexpressing (SLC29A1-OE) cells treated with DMSO or 25 nM IACS-010759 (n=8). Data are from one of three independent experiments. **F.** Tumor growth rates of control (SLC29A1-EV) and SLC29A1-overexpression H460 xenografts. n=14 for each group. **G.** Model of reprogramming of the purine synthesis pathways upon ETC suppression. Unpaired, two-sided t tests (**A-C** and **E**), and a two-way ANOVA test (**F**) were used for the statistical analyses. ****: P < 0.0001; ***: P < 0.001; **: P < 0.01, *: P < 0.05, n.s.: P 0.05. Error bars denote SEM.

NBMPR did not alter proliferation of H460 cells under control conditions, but enhanced the effect of IACS-010759 (Figure 7C) and reduced proliferation in H460 cells lacking UQCRC2 (Figure S7D). SLC29A1 overexpression promoted hypoxanthine uptake and blunted the effect of IACS-010759 on cell proliferation (Figures 7D, 7E and S7E). To test whether purine nucleoside uptake is limiting for H460 growth in vivo, we subcutaneoulsy implanted control and *SLC29A1*-overexpressing cells and assessed xenograft growth. *SLC29A1* overexpression was sufficient to enhance xenograft growth under these conditions (Figure 7F). In line with this, both *SLC29A1* and *SLC29A2* are more highly expressed in human lung adenocarcinoma relative to adjacent lungs, suggesting a role in human cancer (Figure S7F).

## DISCUSSION

Our data indicate that mitochondrial metabolism – specifically, the ability to engage in OXPHOS – dictates the pathway by which cells maintain pools of purines. Genetic or pharmacological blockade of the ETC suppresses de novo purine synthesis, and under these conditions cells require purine uptake and salvage to maximize growth (Figure 7G). We find evidence of alterations in purine metabolites in primary fibroblasts from patients with IEMs affecting mitochondrial function, and in human NSCLCs where the contribution of glucose to the TCA cycle correlates inversely with markers of purine salvage. We also note that cancer patients receiving IACS-010759 in a Phase I clinical trial exhibited elevated purine nucIeotides in the blood^42^, and that defects in mitochondrial DNA replication perturb purine-related metabolites in patients and mice^43^. These findings provide support for the clinical relevance of the findings reported here.

An interesting aspect of the data is that ETC inhibition results not only in a switch from de novo purine synthesis to purine salvage, but an accumulation of the purine monophosphates IMP, GMP, and AMP. Aspartate becomes limiting when the cell’s ability to recycle NADH to NAD^+^ is impaired, particularly under conditions of Complex I dysfunction^27, 28^. Aspartate is required for the conversion of aminoimidazole ribonucleotide to *N*-succinylcarboxamide-5-aminoimidazole ribonucleotide, a step in de novo IMP synthesis. Therefore, the suppression of de novo purine synthesis by ETC blockade is intuitively understandable, but the accumulation of IMP and other purine monophsophates implies that additional factors beyond aspartate limitation contribute to the change in purine metabolism.

The low NAD^+^/NADH ratio induced by ETC dysfunction is a key factor in elevated IMP and hypoxanthine levels, because normalizing this ratio reduces their levels. Although we induced NADH reductive stress by inhibiting Complex I or III of the ETC, the cytosolic rather than mitochondrial NAD^+^/NADH ratio is most relevant to purine monophosphate abundance because Cyto-*Lb*Nox suppresses IMP abundance better than Mito-*Lb*Nox. This effect may be related to the fact that the inosine monophosphate dehydrogenases (IMPDH1/2) are cytosolic NAD^+^-dependent enzymes in the pathway linking IMP to GMP, and their inhibition by a low NAD^+^/NADH ratio is thought to result in IMP accumulation^28^. We note, however, that the high levels of GMP and AMP in addition to IMP imply that additional factors are involved. Other evidence linking purine accumulation to NADH reductive stress includes the observation that expressing the *Escherichia coli* pyridine nucleotide EcSTH in HeLa cells reduces NAD^+^/NADH ratio and increases the abundance of many purines, and that acute ethanol ingestion decreases the NAD^+^/NADH ratio and induces IMP, AMP, and GMP accumulation in the mouse liver^44^. Altogether these observations indicate that NADH reductive stress regulates steady-state purine levels. Our data demonstrate that purine salvage is required for IMP, GMP, and AMP accumulation during Complex I blockade, because HPRT1 knockout eliminates this effect. Furthermore, although the compromised cellular energy state of ETC-inhibited cells may regulate some aspects of purine metabolism, the data imply that the NAD^+^/NADH ratio is the key factor because *Lb*Nox reduces this ratio without restoring OXPHOS.

We speculate that accumulation of purine monophosphates is an adaptive response to help cells cope with compromised ETC function. De novo purine nucleotide synthesis from PRPP to IMP requires ten reactions catalyzed by three multifunctional enzymes. This process is energetically demanding, requires contributions from glutamine, glycine, aspartate and N^10^-formyl-tetrahydrofolate, and is subject to potent feedback inhibition by purine monophosphates^45, 46^. Constitutive purine biosynthesis would be counterproductive and perhaps toxic when the cell’s ability to produce and maintain a favorable energy state and pools of required intermediates is insufficient to complete the purine synthesis pathway^47^. It is therefore interesting that the purine salvage response induced during ETC blockade does not simply replenish purine monophosphate pools, but expands them. This effect may facilitate inhibition of phosphoribosyl pyrophosphate amidotransferase (PPAT)^48^, which catalyzes the committed step of the pathway and is inhibited by multiple purine nucleotides.

The balance between de novo purine synthesis and purine salvage is particularly relevant in cancer, where mitochondrial function is variable and drugs can be used to inhibit either pathway. In some tumors, mutations in mitochondrial enzymes may render cells permanently reliant on purine salvage. Recent data demonstrate that fumarate hydratase (FH)-deficient renal carcinoma cells have suppressed levels of purine synthesis and are highly dependent on purine salvage. The dependence on salvage is relieved by expressing wild-type FH^49^. In NSCLC, both cell lines and tumors have variable mitochondrial activity as assessed by ^13^C labeling of TCA cycle intermediates from [U-^13^C]glucose^18, 21^. We note that H460 cells, which respire well and do not require HPRT1 for growth in culture, nevertheless require this enzyme for maximal growth of subcutaneous xenografts. These findings indicate that dependence on purine salvage can be imposed on respiration-competent cells by environmental factors. Aspartate depletion in xenografts has been reported to compromise the de novo synthesis pathway^50, 51^. Rapidly growing tumors also experience hypoxia, which may further enhance salvage dependence^50, 52, 53^. We emphasize that many xenograft models require some ETC activity for growth^11, 12^, and despite their dependence on HPRT1, the H460 xenografts studied here also displayed growth suppression by IACS-010759. But the environmental factors associated with rapid tumor growth, including hypoxia, may contribute to dependence on purine salvage and explain why NSCLCs tend to over-express *HPRT1*, *SLC29A1* and *SLC29A2*. The fact that over-expressing *SLC29A1* is sufficient to drive xenograft growth argues that access to purine nucleosides or nucleobases is a limiting factor for H460 cell growth in vivo.

## METHODS

### Cell culture

H460, A549, and H157 cells were obtained from the Hamon Cancer Center Collection (University of Texas Southwestern Medical Center) and maintained in RPMI-1640 (Thermo Fisher Scientific, CB-40234) supplemented 10% fetal bovine serum (FBS). 293T and 786-O cells were from American Type Culture Collection (ATCC, CRL-3216; CRL-1932) and maintained in high glucose DMEM with 10% FBS. Patient-derived fibroblasts were cultured in low glucose DMEM (Sigma, D6046) supplemented with 5% heat inactivated FBS. All cells were cultured at 37°C in a humidified atmosphere with 5% CO_2._

### Clinical samples

All subjects were enrolled in a study (NCT02650622) approved by the Institutional Review Board (IRB) at University of Texas Southwestern Medical Center (UTSW). Informed consent was obtained from all patients and their families. Plasma samples were obtained from fresh blood collected in heparinized tubes at the Children’s Medical Center at Dallas. Punch biopsies of the skin for fibroblast culture were obtained from the patient, following the standard procedure for clinical diagnostics.

### Gene deletion and over-expression

To generate *UQCRC2^-/-^* H460 cells, cells were transfected with the PX458 construct^54^, a gift from Feng Zhang (Addgene plasmid #48138; http://n2t.net/addgene:48138; RRID:Addgene_48138) that contains sgRNA against human UQCRC2. Cells with the highest 2-5% GFP signal were selected two to three days post transfection using a flow cytometer (BD FACSAria II). Single cells were cultured in RPMI with 10% FBS, 1mM sodium pyruvate, 100 µg/mL uridine and penicillin/streptomycin. Loss of UQCRC2 protein was validated by western blot. To generate sgScr, sgHPRT1, sgATG5, and sgATG7 H460 cells, the indicated gRNAs were cloned into the LentiCRISPRv2 vector^55^, a gift from Feng Zhang (Addgene plasmid # 52961; http://n2t.net/addgene:52961; RRID:Addgene_52961) and transfected into 293T cells using Lipofectamine 3000 (Thermo Fisher Scientific L3000015) with a 2:1 ratio of psPAX2: pMD2G. The same method was used to generate RPS3-Keima cells using pLENTI_RPS3_Keima construct, a gift from Thomas Tuschl (Addgene plasmid # 127140; http://n2t.net/addgene:127140; RRID:Addgene_127140). For NDI1, LbNOX, and SLC29A1 overexpression, we utilized the PMXS-IRES-Bsd retroviral expression vector. The PMXS-NDI1 vector^56^ was a gift from David Sabatini (Addgene plasmid # 72876; http://n2t.net/addgene:72876; RRID:Addgene_72876). PMXS-Cyto-*Lb*NOX and PMXS-Mito-*Lb*NOX were a gift from Kivanc Birsoy. To generate stable gene-overexpressing H460 cell lines, the indicated PMXS constructs were transfected into 293T cells using Lipofectamine with a 2:1 ratio of Gag-Pol:VSVG. After 48 hours of transfection, medium containing viral particles was harvested and filtered using 0.45µm membranes and immediately used to culture H460 cells in the presence of 4 µg/mL polybrene (Sigma, TR-1003-G). After 24 hours, cells were subjected to 2 µg/mL puromycin (Thermo Fisher Scientific, NC9138068) or 10 µg/mL blasticidin (Thermo Fisher Scientific, NC1366670) selection until all the uninfected cells died. DNA oligos were purchased from IDT and contained the following sequences:

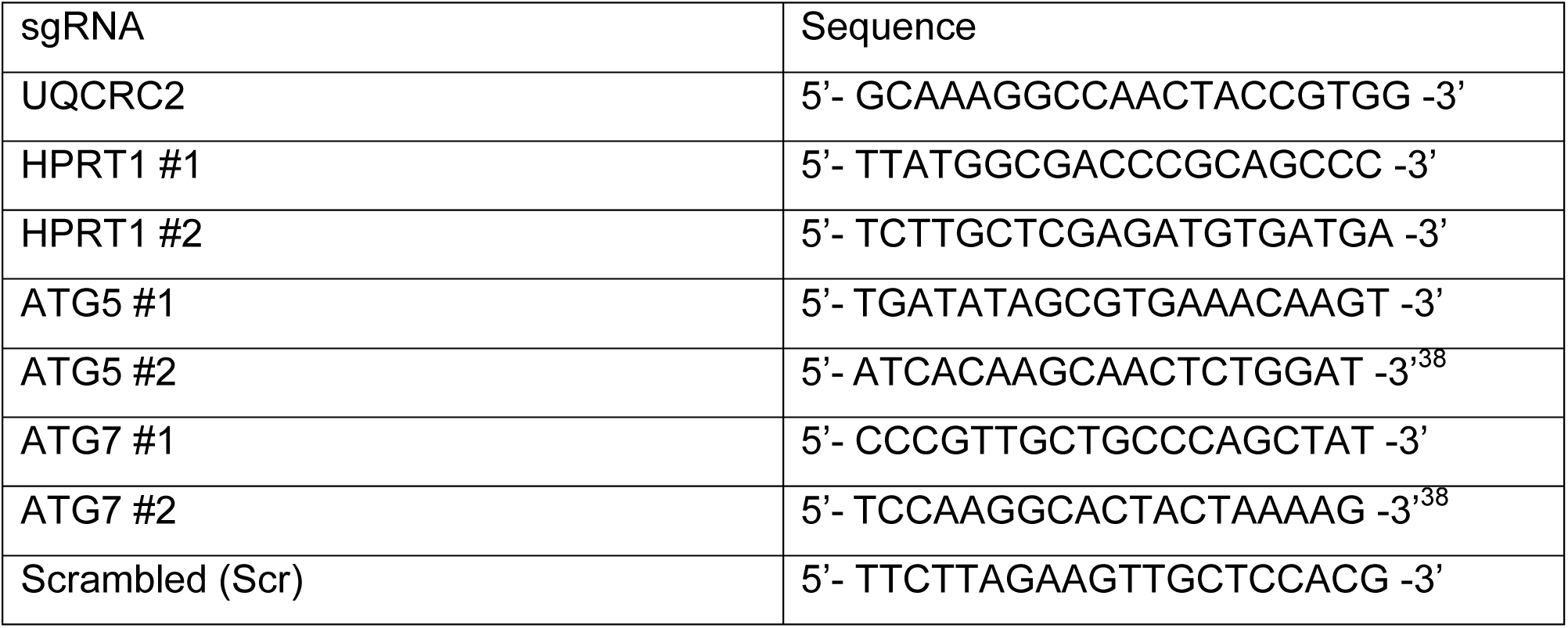

### NAD^+^ and NADH quantitation

Qualitative analysis of NAD^+^ and NADH was performed as previously described^57^ on a QExactive HF-X mass spectrometer (Thermo Scientific, Bremen, Germany). We perfomed quantitative analysis of NAD^+^ and NADH according to our previous protocol^58^ on a 6500+ mass spectrometer (AB Sciex, Framingham, MA). To prepare quantitative samples, cells were washed with saline and extracted with 40:20:20 acetonitrile:methanol:water (v/v) with 0.1 M formic acid and then neutralized with 15% ammonium bicarbonate (w/v). A ^15^N_5_-AMP internal standard was added to each extract at the final concentration of 100 nM. Samples were run the same day to minimize oxidation of the analytes of interest.

All cellular extracts were analyzed against an 8-point standard curve ranging from 5 nM to 1000 nM. All standard curves had R^2^ values greater than or equal to 0.98 with greater than 6 calibrators having accuracies within 20% of their known concentration.

### Targeted metabolomics

To extract metabolites, cells were rinsed with ice-cold saline twice and quenched by 80% cold methanol. Cells were incubated at –80°C for at least 20 minutes and then scraped. For tumor samples, the tissues were thawed and homogenized in the cold 80% methanol using plastic pestles (Thermo Fisher Scientific, 12141364). Samples were subjected to three freeze-thaw cycles in liquid nitrogen and a 37°C water bath. Afterwards, the samples were vortexed for 1 minute and spun down at 4°C at the 20,160 x g for 15 minutes. The supernatants were transferred into fresh Eppendorf tubes and dried in a SpeedVac concentrator overnight.

Metabolite abundance was analyzed using multiple mass spectrometers. For analysis on a Q-TOF mass spectrometer, dried metabolites were reconstituted in 0.1% formic acid in analytical-grade water and vortexed for 1 minute before spinning at 4°C at 20,160 x g for 15 minutes. The supernatants were transferred to auto-sampler vials. Data acquisition was performed by reverse-phase chromatography on a 1290 UHPLC liquid chromatography (LC) system interfaced to a high-resolution mass spectrometry (HRMS) 6550 iFunnel Q-TOF mass spectrometer (MS) (Agilent Technologies, CA). The MS was operated in both positive and negative (ESI+ and ESI-) modes. Analytes were separated on an Acquity UPLC® HSS T3 column (1.8 µm, 2.1 x 150 mm, Waters, MA). The column was kept at room temperature. Mobile phase A composition was 0.1% formic acid in water and mobile phase B composition was 0.1% formic acid in 100% ACN. The LC gradient was 0 min: 1% B; 5 min: 5% B; 15 min: 99%; 23 min: 99%; 24 min: 1%; 25 min: 1%. The flow rate was 250 µL min^-1^. The sample injection volume was 5 µL. ESI source conditions were set as follows: dry gas temperature 225 °C and flow 18 L min^-1^, fragmentor voltage 175 V, sheath gas temperature 350 °C and flow 12 L min^-1^, nozzle voltage 500 V, and capillary voltage +3500 V in positive mode and −3500 V in negative. The instrument was set to acquire over the full *m/z* range of 40–1700 in both modes, with the MS acquisition rate of 1 spectrum s^-1^ in profile format.

Raw data files (.d) were processed using Profinder B.08.00 SP3 software (Agilent Technologies, CA) with an in-house database containing retention time and accurate mass information on 600 standards from Mass Spectrometry Metabolite Library (IROA Technologies, MA) which was created under the same analysis conditions. The in-house database matching parameters were: mass tolerance 10 ppm; retention time tolerance 0.5 min. Peak integration result was manually curated in Profinder for improved consistency and exported as a spreadsheet (.csv).

Samples prepared for analysis on a Q-Exactive were reconstituted in 80% acetonitrile and centrifuged at 4°C at 20,160 x g to remove insoluable material. Chromatographic separation of metabolites was carried out on a Vanquish UHPLC system equipped with a ZIC-pHILIC column (Millipore-Sigma, Burlington, MA) as previously described ^57, 59, 60^. Extracted ion chromatograms (XICs) were generated with a mass tolerance of 5 ppm and integrated for relative quantitation. Identities of analytes were confirmed with purified standards and product ion spectra.

Principal component analyses and metabolite set enrichment analyses were conducted using MetaboAnalyst 5.0^61^.

### Stable isotope tracing

For glucose tracing, cells were cultured in base RPMI medium (Sigma, R1383-L) supplemented with 11 mM [U-^13^C]glucose (Cambridge Isotope Laboratories, CLM481-0.25) or [1,2-^13^C]glucose (Cambridge Isotope Laboratories, CLM-504-0.5) and 10% dialyzed FBS (Gemini Bio-Products, 100108). For glutamine tracing, cells were cultured in glutamine-free RPMI medium (Sigma, R0883) supplemented with 2 mM [amide-^15^N]glutamine (Cambridge Isotope Laboratories, NLM-557-1) and 10% dialyzed FBS. For hypoxanthine tracing, cells were cultured in RPMI medium containing 10% dialyzed FBS and 10 µM [^15^N_4_]hypoxanthine (Cambridge Isotope Laboratories, NLM-8500-0.1). For all drug-treated samples, cells were exposed to the drug during tracing. The metabolites were extracted as described in the targeted metabolomics method. For analysis by Q-TOF, data acquisition was performed and analyzed according to the methods described above.

For analysis on the Q-Exactive, tSIM methods were used to increase the signal of isotopically-labeled intermediates of the purine biosynthetic pathway and pentose phosphate pathway. Both ^15^N and ^13^C nuclei were analysed using this approach. Quadrupole isolation windows for individual analytes were set to capture all relevant nuclei to calculate fractional enrichment values. We performed analysis of isotopologues according to our previously reported method^62^. Natural isotope abundances were corrected using a customized R script, which can be found at the GitHub repository (https://github.com/wencgu/nac). The script was written by adapting the AccuCor algorithm^63^.

For targeted analysis of purines and hypoxanthine tracing using AB SCIEX QTRAP 5500 LC/triple quadrupole MS (Apllied Biosystems SCIEX), metabolites were reconstituted in 0.1% formic acid in analytical water, vortexed, and spun down to remove insoluble material before being loaded onto the instrument as previously described^64^. Using a Nexera Ultra-High-Performance Liquid Chromatograph system (Shimadzu Corporation), we achieved separation on a Waters Symmetry C18 column (150 × 2.1 mm, 3.5um) with 0.1 % formic acid in mobile phase A (H_2_O) and mobile phase B (acetonitrile) in a flow rate at 0.4 mL with injection volume at 10 µL. The gradient elution is 0-5 min, 0-30% B; 5-6 min, 30-100% B; 6-8 min, 100% B; 8-9 min, 100-0% B; 9-10 min, 12% B. Chromatogram review and peak area integration were performed using MultiQuant (version 2.1, Applied Biosystems SCIEX). The MRMs used are listed as follows: AMP Q1/Q3 (348/136 (m+0), 352/140 (m+4), CE: 30); IMP Q1/Q3 (349/137 (m+0), 353/141 (m+4), CE: 22); GMP Q1/Q3 (364/152 (m+0), 368/156 (m+4), CE: 18); Hypoxanthine Q1/Q3 (137/119 (m+0), 141/123 (m+4), CE: 27) or (137/110 (m+0), 141/113 (m+4), CE: 27).

### HPRT1 enzymatic activity analysis

HPRT1 enzyme activity was measured using the Precise HPRT1 assay kit (Novo CIB, K0709-01-2) according to manufacturer’s instructions. In brief, H460 cells were seeded in 10 cm plates and treated with DMSO or 25 nM IACS-010759 for 24 hours. Cells were rinsed once with PBS, scraped, and lysed in the ice-cold lysis buffer containing 10 mM Tris-HCl pH 7.4, 150 mM NaCl, 1% NP-40, 1 mM EDTA followed by centrifugation at 18,000 x g for 10 min at 4 °C. Each enzymatic reaction contains 5 µL sample or positive control (human recombinant HPRT enzyme) and 200 µL reaction mixture containing DTT (cofactor 1), NAD (cofactor 2) and bacterial IMPDH in the absence (blank) or presence of 2 mM PRPP (enzyme reaction). The reaction was performed at 37 °C. The absorbance at 340 nm was recorded at 2-minutes intervals for 120 minutes. HPRT1 protein abundance in each sample was assessed by immunoblotting using antibody against HPRT1. Image J was used to quantify the HPRT1 band intensity. HPRT1 catalytic rate was normalized to the protein abundance in each group with the DMSO control samples given a value of 1.

### Glucose uptake and lactate secretion assay

Cells were seeded in 6 cm plates. The glucose uptake and lactate secretion analysis was started when cells reached 90% confluence. Cells were washed with PBS once. 2 mL medium containing 25 nM IACS-010759 or equal volume of DMSO was added into the plates for 6 hours. Medium from each plate was collected and spun down at 20,160 xg at 4°C. 1 mL supernatant of each sample was transferred to a fresh eppendorf tube and loaded in a NOVA instrument to measure glucose and lactate levels. Three or four tubes containing medium but no cells were used as blanks to calculate the amount of glucose and lactate taken up or secreted by cells. The cell number was counted from each plate to calculate the rate of glucose uptake and lactate secretion per cell.

### Cell growth analysis

Cells were seeded in flat, clear bottom 96 well plates. Cells were stained with 5 µg/mL Hoechst (Thermo Fisher Scientific, 62249) and 1 µg/mL Propidium Iodide (PI) (Thermo Fisher Scientific, P3566) in PBS for at least 15 minutes at 37°C and then subjected to cell counting using a Celigo Imaging Cytometer. Live cells were calculated as the total number of Hoechst-positive cells minus the number of PI-positive cells. Proliferation rate was calculated as previously described^23^. Chemicals added to the medium for the growth assay were: IACS-010759 (25nM, ChemieTek, CT-IACS107), pyruvate (1mM, Sigma-Aldrich, S8636), uridine, AKB (1mM, Sigma-Aldrich, K0875-5G), inosine (50µM, Millipore Sigma, I4125), guanosine (50µM, Millipore Sigma, G6752), adenosine (50µM, Millipore Sigma, A9251), and NBMPR (50µM, N2255, Millipore Sigma).

### Immunoblotting

Cells were washed with PBS and then lysed in RIPA buffer (Boston BioProducts, BP-115) containing proteinase and phosphatase inhibitors (Thermo Fisher Scientific, 78444). Samples were centrifuged at 4°C at 20,160 x g for 10 minutes and supernatants were collected for protein measurement using the DC Protein Assay Kit (Bio-Rad, 5000111). Equal amounts of protein were loaded to run the gels (Thermo Fisher Scientific, NP0323BOX) and then transferred to PVDF membrane (Thermo Fisher Scientific, 88518). Membranes were dipped in methanol for 20 seconds and then rinsed with DI water. The air-dried membrane was incubated with primary antibodies in filtered PBS containing 5% BSA and 0.1% Tween-20 (PBST) at 4°C overnight. The membranes were washed with PBS 3 times for 5 minutes and then incubated with horseradish peroxidase conjugated secondary antibody (Cell Signaling Technology, 7074, 7076) in 5% non-fat milk in PBST at room temperature for 1 hour. Membranes were washed 5 times for 5 minutes with PBS at room temperature and then exposed to Pierce ECL (Thermo Fisher Scientific, PI32106) for minutes. Signals were detected using either Amersham imagequant 800 or films in the dark room. Antibodies used for western blots are: Vinculin (Proteintech, 26520-1-AP), UQCRC2 (Abcam, ab14745), HPRT1 (Santa Cruz Biotechnology, sc-376938), SLC29A1 (Abcam, ab182023), p70 S6 Kinase (S6K) (Cell Signaling Technology, 9202S), Phospho-p70 S6 Kinase P-S6K (Thr389) (Cell Signaling Technology, 9234S), 4E-BP1 (Cell Signaling Technology, 9644S), Phospho-4E-BP1 (Ser65) (Cell Signaling Technology, 9451S), GAPDH (Cell Signaling Technology, 2118S), RPL26 (Bethyl Laboratories, A300-685A-T), Keima (MBL International, M1823B), LC3B (Sigma-Aldrich, L8918), p62 (Cell Signaling Technology, 88588S), ATG5 (Cell Signaling Technology, 9980S), ATG7 (Cell Signaling Technology, 8558T).

### Xenograft studies in mice

All mouse experiments complied with relevant ethical regulations and were performed according to protocols approved by the Institutional Animal Care and Use Committee at the University of Texas Southwestern Medical Center (protocols 2016-101360 and 2016-101694). H460 cells were suspended in serum-free RPMI medium and mixed with Matrigel (Thermo Fisher Scientific, CB-40234) at 1:1 volume ratio. One million cells were subcutaneously injected into the right flank of NOD.Cg-*Psrkdcscid Il2rgtm1Wjl*/SzJ (NSG) mice. Mice were randomized for IACS-010759 treatments. Mice were administrated with IACS-010759 daily through oral gavage (5 or 10mg/kg body mass in 100 µL of 0.5% methylcellulose and 4% DMSO)^26, 30^. For tumor metabolomics and glutamine infusion experiments, mice were dosed with (10mg/kg) IACS-010759 for 5 days prior to the infusion as previously described^30^. After the infusion, tumors were collected and snap frozen for later metabolite extraction and LC-MS analyses. For tumor growth analyses, mice were administered 5 mg/kg IACS-010759 until the day before sacrifice. Two orthogonal measurements of tumor diameter were collected every other day and tumor volume was calculated using the formula V = (L_1_x(L_2_^2^))/2.

### [amide-^15^N]glutamine infusion

Mice were anesthetized using (30 mg/mL) ketamine/xylazine mix (30 µL/g). 25-gauge catheters were placed in the lateral tail vein under anesthesia. The total dose of glutamine was 1.725 g/kg dissolved in 1.5 mL saline. Isotope infusions started with 150 µL/minute bolus for 1 minute followed by continuous infusion at rate of 150 µL/hour for 4 hours. Upon termination of the infusions, animals were euthanized immediately and tumors were collected and snap frozen in liquid nitrogen.

### Seahorse XFe96 Respirometry

An XFe96 Extracellular Flux Analyzer (Agilent Technologies) was used to measure oxygen consumption rate. In brief, 20,000 cells per well were seeded and simultaneously treated with 25 nM IACS-010759. After 16 to 20 hours, cells were washed three times with seahorse medium (Agilent Technologies, 102353) containing 2 mM glutamine, 1 mM pyruvate, 10 mM glucose and pen/strep (pH 7.4) and incubated in a CO_2_-free incubator at 37°C for at least 30 minutes prior to loading into the instrument. Final concentrations for oligomycin A, carbonyl cyanide m-chlorophenylhydrazone (CCCP), and rotenone are 2 µM, 1 µM, and 2 µM respectively. After the assay, cells were counted using Celigo Image Cytometer (see method: cell growth analysis) to normalize oxygen consumption rate.

### Immunofluorescence and confocal microscopy

Coverslips coated with 10 µg/mL fibronectin (Sigma-Aldrich, F1141-5MG) for 1 hour at 37°C and rinsed once with PBS. Cells were immediately seeded on the coverslips. Cells were fixed the next day with fresh warm 4% paraformaldehyde (PFA) solution in PBS for 15 minutes followed by permeabilization using 0.1% (v/v) Triton X-100 in PBS at room temperature for 10 minutes. Cells were then blocked in filtered PBS containing 1% BSA for at least 30 minutes at room temperature before incubation with primary antibodies against FLAG (1:200, F1804, Sigma-Aldrich) and HSP60 (1:500, 12165S, CST) for 1 hour at room temperature. Cells were washed 3 times for 5 minutes with PBS and then incubated with secondary antibodies (Alexa fluorophores 488 and 555, Invitrogen) for 1 hour in dark at room temperature. Coverslips were washed with PBS 3 times for 5 minutes and Mili-Q water once before being mounted on slides by Profade-Antifade (P36935, Invitrogen) overnight in dark. Cells were imaged using Zeiss LSM 880 Confocal Laser Scanning Microscope with Z-stacks acquired. All representative images were processed using Image J.

### RNA-seq

RNA was extracted using Trizol (Thermo Fisher Scientific, 15596018) and an RNeasy Mini Kit (Qiagen, 74106). Qubit fluorometer and the Invitrogen Qubit RNA High Sensitivity kit (Invitrogen, Q32852) were used to measure total RNA levels. RNA-seq libraries were prepared using the NEBNext Ultra II directional RNA library prep kit with the NEBNext Poly(A) mRNA magnetic isolation module (New England Biolabs, E7490L, E7760L) according to manufacturer’s instructions. Libraries were stranded using standard N.E.B indices according to manufacturer’s instructions (New England Biolabs, E7730L, E7335L, E7500L). Sequencing reads from all RNA-seq experiments were aligned to hg19 reference genome by STAR v. 2.5.2b^65^ with the following parameters: –– runThreadN 28 ––outSAMtype BAM SortedByCoordinate ––outFilterMultimapNmax 1 –– outWigStrand Unstranded ––quantMode TranscriptomeSAM. Output BAM files were converted to BED format using the “bamtobed” command from BEDtools v.2.29.2 [https://bedtools.readthedocs.io/en/latest/]. BED files were then converted to a normalized wiggle file using a custom python script. Normalized wiggle files were then converted to bigwig format using wigToBigWig with “-clip” parameter. Read counts were derived using HTSeq^66^ with parameter “-s no” and 1 additional read count was added to each gene for each independent sample prior to downstream analyses. Differentially expressed genes were identified by DESeq2 (fold change ≥ 1.5, FDR-adjusted P value ≤ 0.05)^67^. Fragments Per Kilobase Of Exon Per Million Fragments Mapped (FPKM) value of genes were calculated by normalizing the gene length and sequencing depth.

## STATISTICAL ANALYSIS

Figures were prepared and statistics were calculated using GraphPad PRISM. Unless otherwise indicated in the figure legends, statistical significance was calculated using an unpaired, two-tailed student’s t-test with 95% confidence intervals. *P < 0.05; **P < 0.01; ***P < 0.001; ****P < 0.0001; NS, not significant (P > 0.05). Statistical details can also be found in the figure legends for each figure.

## ACKNOWLEDGEMENTS

We thank Gerta Hoxhaj for advice on ribophagy analysis, and we thank members of the DeBerardinis laboratory and Aron B. Jaffe for critically assessing the work. This article is subject to HHMI’s Open Access to Publications policy. HHMI lab heads have previously granted a nonexclusive CC BY4.0 license to the public and a sublicensable license to HHMI in their research articles. Pursuant to those licenses, the author-accepted manuscript of this article can be made freely available under a CC BY4.0 license immediately upon publication. This research was supported by the Howard Hughes Medical Institute Investigator’s Program (R.J.D) and grants from the National Cancer Institute (R35CA22044901, P50CA070907 and P50CA196516). D.B. was supported by grants from the N.I.H (F31CA239330, T32GM008203, TL1TR001104).

## AUTHOR CONTRIBUTIONS

R.J.D., M.N., and Z.W., conceived and designed the study. R.J.D. and M.N. recruited patients. Z.W. designed and conducted experiments; and analyzed the data. T.P.M., H.S.V., F.C. and M.S.M.S. designed, optimized and validated the metabolomics methods. R.J.D. and Z.W. interpreted the data and wrote the manuscript. D.B., R.C.H, C.P., F.C., H.S.V., H.C., P.T.N., M.S.M.S., D.D., B.F., L.C., Y.Z., W.G., B.K., B.B., S.K., Y.Z., K.C.O., J.G.B., M.N., L.G.Z. conducted the experiments or analyzed the data.

## DATA AND MATERIALS AVAILABILITY

Requests for the data should be submitted to the corresponding author.

**Supplemental Figure 1**

**A-B.** Glucose uptake (**A**) and lactate secretion (**B**) in H460 cells treated with DMSO or 25 nM IACS-010759 for 6 hours. Data are means from three replicates.

**C.** Principal component analysis of metabolomic profiles in H460 cells treated with DMSO or 25 nM IACS-010759 for 24 hours.

**D.** Relative abundance of the indicated metabolites in H460 cells treated with DMSO or 25 nM IACS-010759. Data are means from three replicates.

**E.** Oxygen consumption rates in the indicated cell lines pre-treated with DMSO or 25 nM IACS-010759 for 24 hours. OA: oligomycin A; R: rotenone. Data are from one of three independent experiments.

**F.** Relative purine nucleotide abundance in the indicated cell lines treated with DMSO or 25 nM IACS-010759 for 24 hours. Data are means from three replicates.

**G.** Principal component analysis of metabolomic profiles in H460 xenografts treated with vehicle or IACS-010759 for five days.

**H.** Metabolite set enrichment analysis comparing vehicle and IACS-010759-treated H460 xenografts.

**I.** Relative abundance of the indicated metabolites in H460 xenografts treated with vehicle or IACS-010759 for five days. Vehicle (n=10), IACS-010759 (n=8).

Unpaired, two-sided t tests were used for the statistical analyses. ****: P < 0.0001; ***: P < 0.001; **: P < 0.01, *: P < 0.05. Error bars denote SEM.

**Supplemental Figure 2**

**A.** Glucose uptake and lactate secretion in H460 cells treated with DMSO or 25 nM IACS-0107596 for 6 hours. Data are means from three replicates.

**B.** Growth rates of control and NDI1-expressing cells cultured in glucose or galactose medium treated with DMSO or 25 nM IACS-010759 (n=6). Data are from one of three independent experiments.

**C-D.** Heatmaps showing metabolomic profiles (**C**) and purine metabolite abundance (**D**) in control and NDI1-expressing cells treated with DMSO or 25 nM IACS-010759 for 24 hours.

**E.** Metabolite set enrichment analysis comparing IACS-010759-treated control and NDI-expressing H460 cells.

Unpaired, two-sided t tests were used for the statistical analyses. ***: P < 0.001; n.s.: P > 0.05. Error bars denote SEM.

**Supplemental Figure 3**

**A.** Western blot validating deletion of UQCRC2. Vinculin is the loading control.

**B.** Oxygen consumption rates in WT and *UQCRC2^-/-^* H460 cells. OA: oligomycin A; AA; antimycin A. Data are from one of three independent experiments.

**C.** Growth rates of WT and *UQCRC2^-/-^* H460 cells. Data are from one of three independent experiments.

**D.** Principal component analysis of metabolomic profiles in WT and *UQCRC2^-/-^* H460 cells.

**E.** Heatmap showing metabolomic profiles in WT and *UQCRC2^-/-^* H460 cells.

**F.** Growth rates of *UQCRC2^-/-^* H460 cells expressing empty vector (EV), Mito-LbNOX, or Cyto-LbNOX (n=10). Data are from one of three independent experiments.

**G.** Relative abundance of aspartate in WT H460 cells and *UQCRC2^-/-^* H460 cells expressing empty vector (EV), Mito-LbNOX or Cyto-LbNOX. Data are means from three replicates.

**H.** Heatmap showing metabolomic profiles in WT H460 cells and *UQCRC2^-/-^* H460 cells expressing empty vector (EV), Mito-*Lb*NOX or Cyto-*Lb*NOX.

Unpaired, two-sided t tests were used for the statistical analyses. ****: P < 0.0001; ***: P < 0.001. Error bars denote SEM.

**Supplemental Figure 4**

**A.** Heatmap showing mRNA levels (FPKM) of purine pathway-associated genes in control and NDI1-expressing H460 cells treated with DMSO or 25 nM IACS-010759 for 24 hours.

**B.** Isotopologue fractions in GTP and ATP after 6 hours of culture with [U-^13^C]glucose in control and NDI1-expressing H460 cells treated with DMSO or 25 nM IACS-010759 for 24 hours. Data are means from three replicates.

**C.** Time-dependent m+4 enrichment in GMP and AMP during culture with [^15^N_4_]hypoxanthine in H460 cells pre-treated with DMSO or 25 nM IACS-010759 for 24 hours. Data are means from three replicates.

**D.** Western blot validating deletion of HPRT1. Vinculin is the loading control.

**E.** Fractional enrichment of m+4 hypoxanthine after 6 hours of culture with [^15^N_4_]hypoxanthine, following 24 hours of pre-treatment with DMSO or 25 nM IACS-010759, in control (sgScr) or HPRT1-depleted (sgHPRT1) H460 cells. Data are means from three replicates.

**F.** Relative HPRT1 enzymatic activity in lysates from H460 cells treated with DMSO or 25 nM IACS-010759 for 24 hours. Data are means from three replicates.

**G.** Relative abundance of R5P in H460 cells treated with DMSO or 25 nM IACS-010759 for 24 hours. Data are means from three replicates.

**H.** Schematic illustrating R5P labeling from [U-^13^C]glucose.

**I.** Relative abundance of R5P during [U-^13^C]glucose tracing in H460 cells pre-treated with DMSO or 25 nM IACS-010759 for 24 hours. Data are means from three replicates.

**J.** Time-dependent m+5 enrichment in R5P from [U-^13^C]glucose in H460 cells pre-treated with DMSO or 25 nM IACS-010759 for 24 hours. Data are means from three replicates.

**K.** Time-dependent fractional enrichment of m+6 6-PG and m+7 S7P during [U-^13^C]glucose tracing in H460 cells pre-treated with DMSO or 25 nM IACS-010759 for 24 hours. Data are means from three replicates.

Unpaired, two-sided t tests (**E-G**) and two-way ANOVA test (**C, I-K**) were used for the statistical analyses. ****: P < 0.0001; ***: P < 0.001, **: P < 0.01, *: P < 0.05; n.s.: P > 0.05. Error bars denote SEM.

**Supplemental Figure 5**

**A.** Correlation between m+2 malate and m+2 glutamate in human NSCLC tumors subjected to intra-operative infusion with [U-^13^C]glucose.

**B.** Fractional enrichment of m+2 glutamate and relative IMP abundance in human NSCLCs with high or low glutamate labeling. The analysis used tumors in the top or bottom 25% of glutamate m+2 labeling (n=7 of each).

**C.** Fractional enrichment of m+2 glutamate and *HPRT1* mRNA levels in human NSCLCs with high or low glutamate labeling. The analysis used tumors in the top or bottom 25% of glutamate m+2 labeling (n=6 of each).

**Supplemental Figure 6**

**A.** Western blot assessing mTORC1 signaling in H460 cells treated with 25 nM IACS-010759. GAPDH and RPL26 are loading controls.

**B.** Western blot assessing mTORC1 signaling in control and NDI1-expressing H460 cells treated with DMSO or IACS-010759 for 24 hours. GAPDH is the loading control.

**C.** Western blot validating RPS3-Keima expression in H460 cells. Calnexin is the loading control.

**D.** Western blot assessing ribophagy and overall macroautophagy in RPS3-Keima-expressing H460 cells treated with DMSO or the indicated compound(s) for 24 hours. GAPDH is the loading control.

**E.** Western blot validating ATG5 and ATG7 deletion. Vinculin and GAPDH are loading controls.

**F.** Relative abundance of the indicated purine nucleotides after 24 hours of DMSO or 25 nM IACS-010759 treatment in control (sgScr), ATG5-deficient (sgATG5), and ATG7-deficient (sgATG7) H460 cells. Data are means from three replicates.

Error bars denote SEM.

**Supplemental Figure 7**

**A.** Growth rates of H460 cells treated with 25 nM IACS-010759 and cultured in medium containing or lacking a mixture of 50 µM adenosine, inosine, and guanosine nucleosides (n=10). Data are from one of three independent experiments.

**B.** Heatmap showing RNA levels of SLC28A and SLC29A family transporters in parental, control, and NDI1-expressing H460 cells with or without 24 hours of IACS-010759 treatment.

**C.** Relative abundance of m+4 hypoxanthine in H460 cells after 6 hours of culture with [^15^N_4_]hypoxanthine, and following a 24-hours pre-treatment with the indicated compounds. Data are means from four replicates.

**D.** Growth rates of *UQCRC2^-/-^*cells treated with DMSO or 50 µM NBMPR (n=10). Data are from one of three independent experiments.

**E.** Relative abundance of m+4 hypoxanthine in conditioned medium collected after incubating control (SLC29A1-EV) and SLC29A1-overexpressing (SLC29A1-OE) H460 cells with [^15^N_4_]hypoxanthine for 2 hours. Data are means from three replicates.

**F.** *SLC29A1* and *SLC29A2* RNA levels in human lung adenocarcinoma (LUAD). N: nonmalignant lung; T: tumors. Data and statistics were generated using TIMER 2.0^68, 69^. Unpaired, two-sided t tests (**A** and **C-E**) were used for the statistical analyses. ****: P < 0.0001; ***: P < 0.001; **: P < 0.01; *: P < 0.05. Error bars denote SEM.

