## Supplemental figures for "Electron transport chain inhibition increases cellular dependence on purine transport and salvage"

### Supplemental Figure 1 (Related to Figure 2)

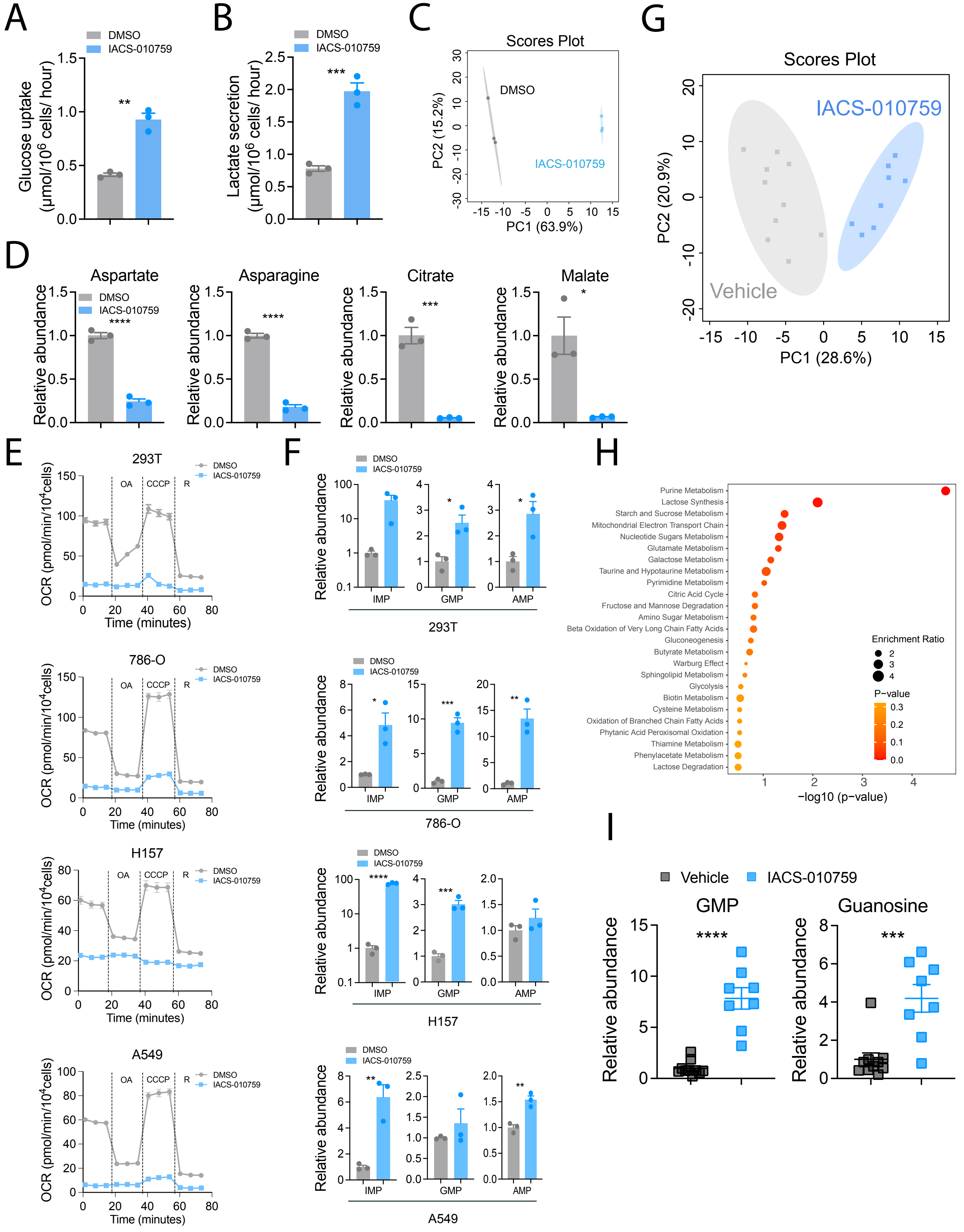

### Supplemental Figure 2 (Related to Figure 2)

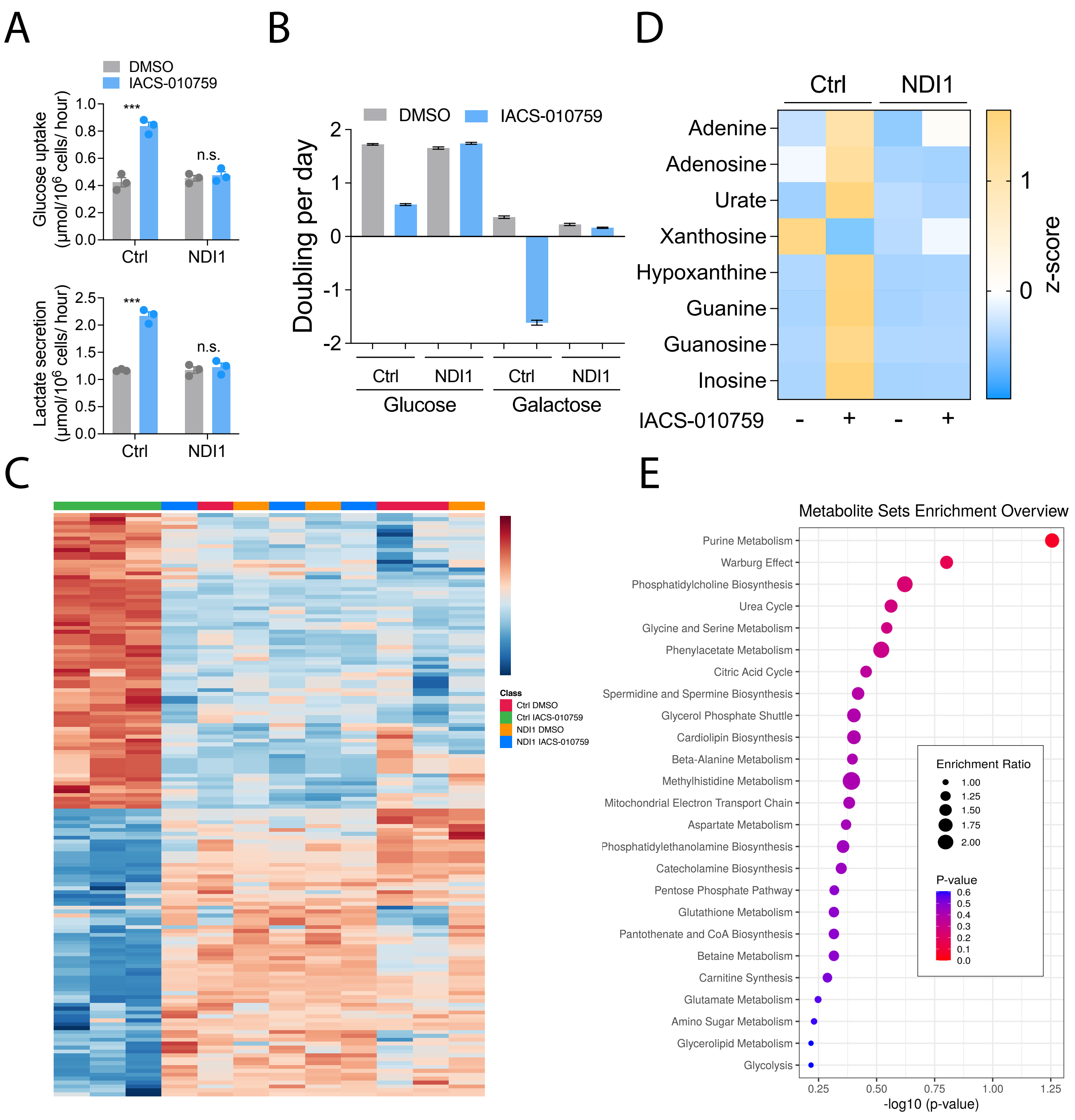

### Supplemental Figure 3 (Related to Figure 3)

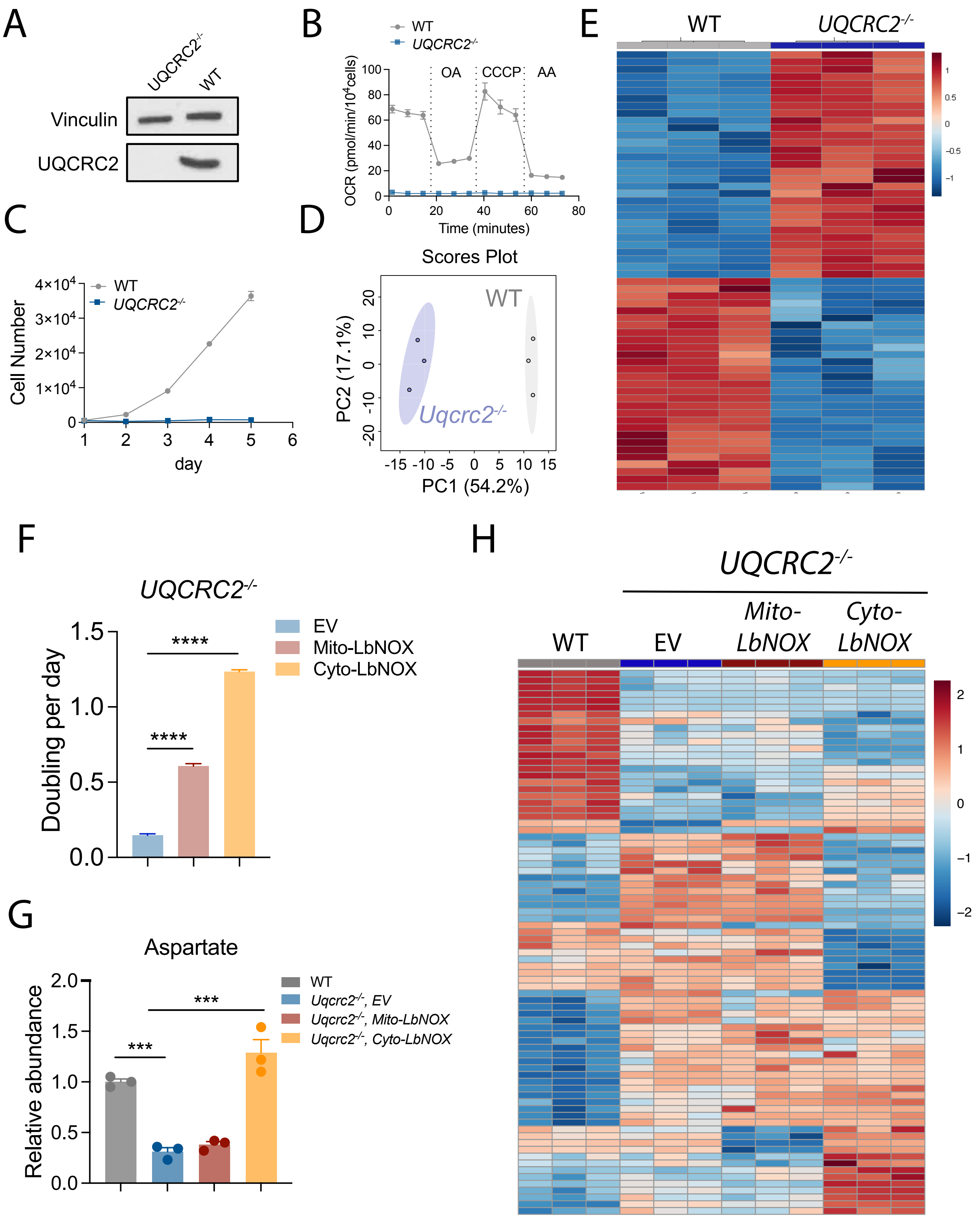

### Supplemental Figure 4 (Related to Figure 5)

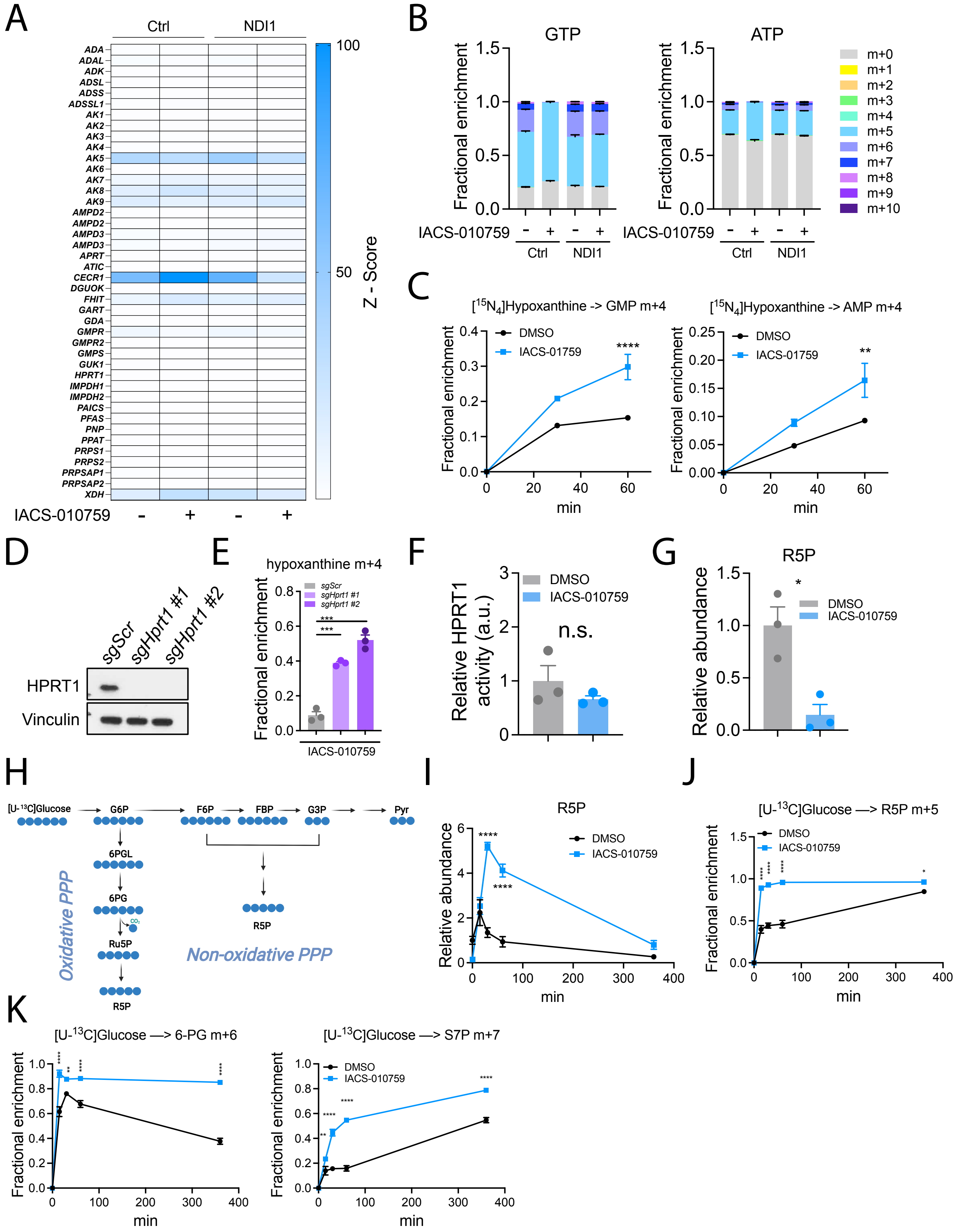

### Supplemental Figure 5 (Related to Figure 6)

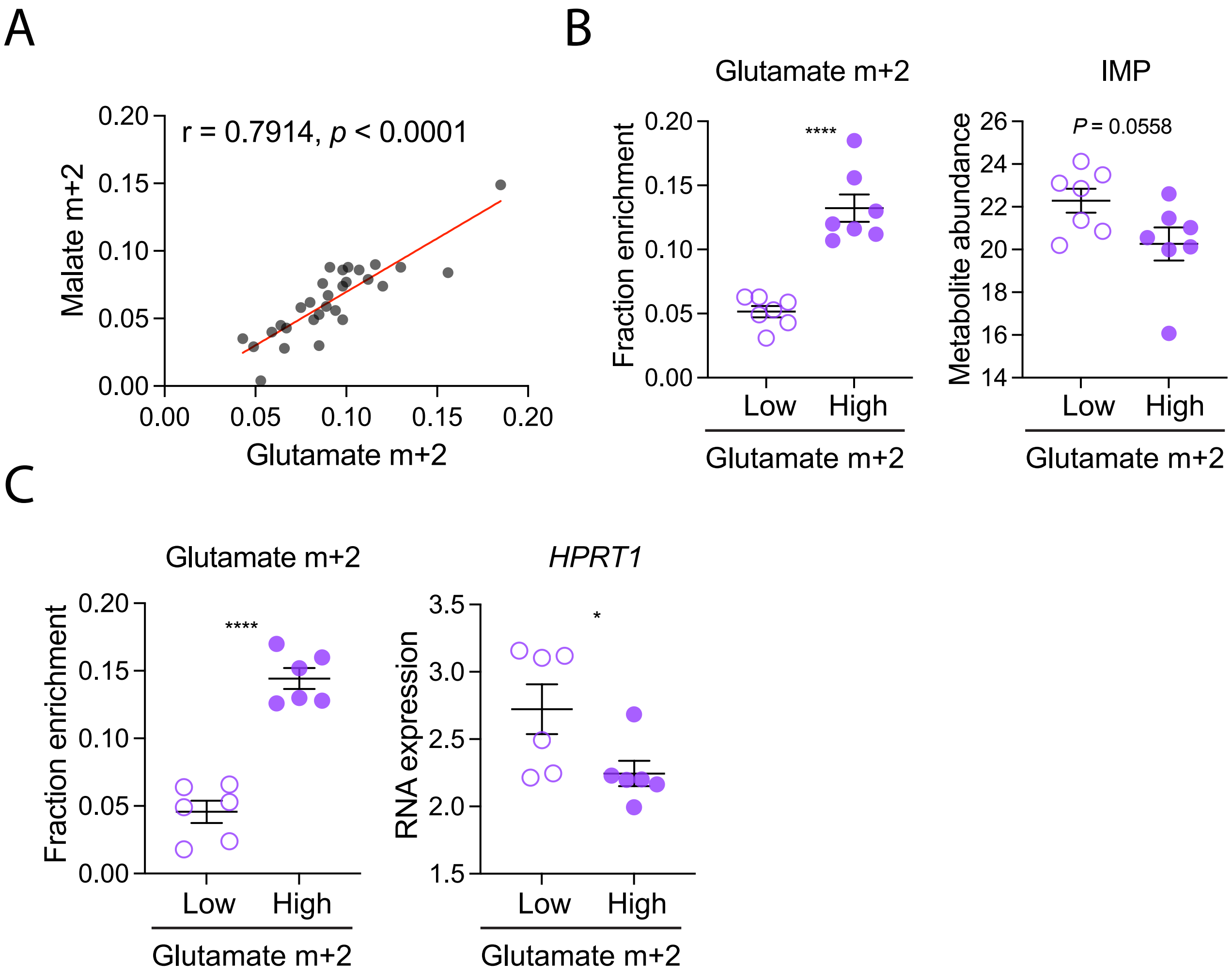

### Supplemental Figure 6 (Related to Figure 7)

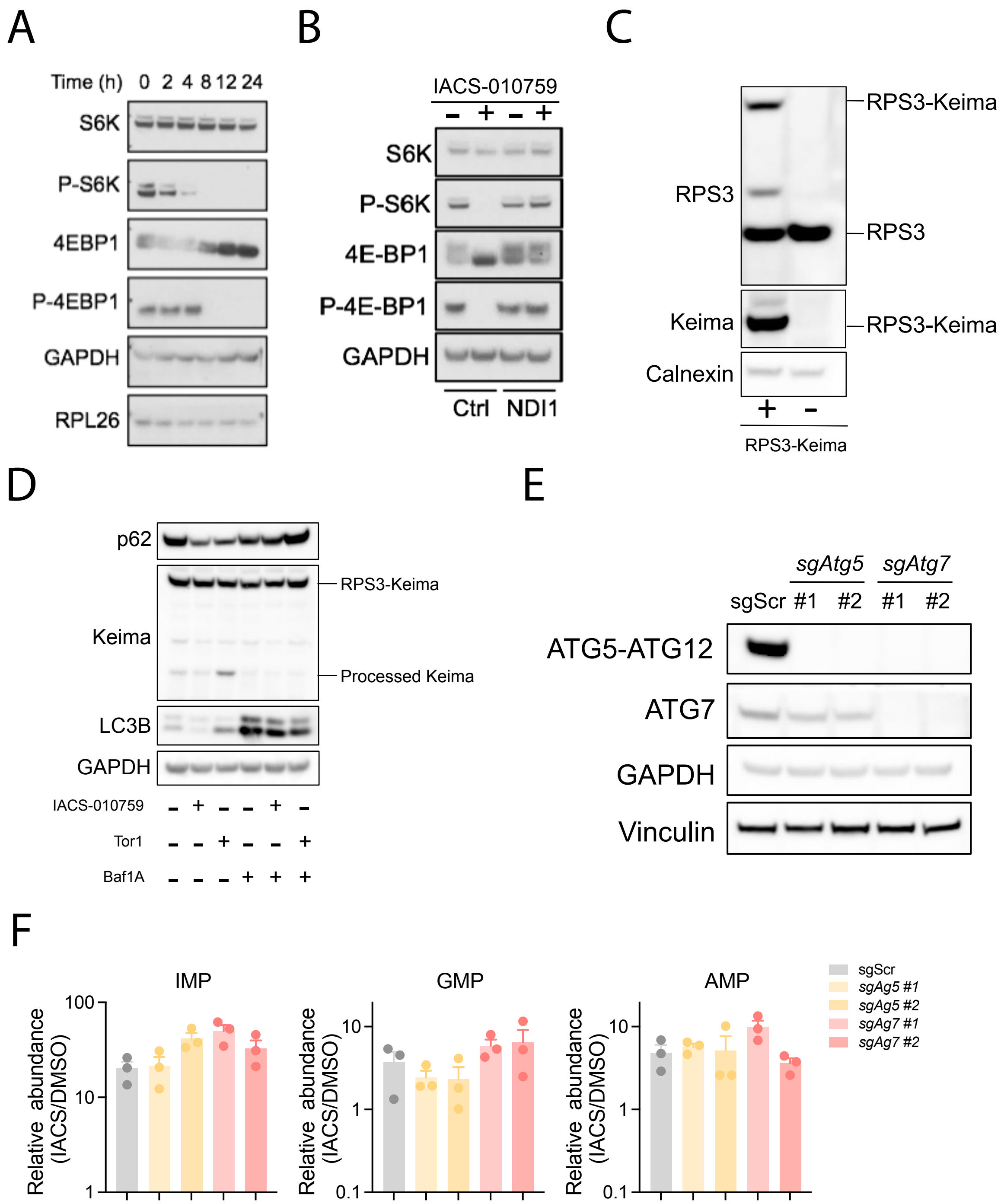

### Supplemental Figure 7 (Related to Figure 7)

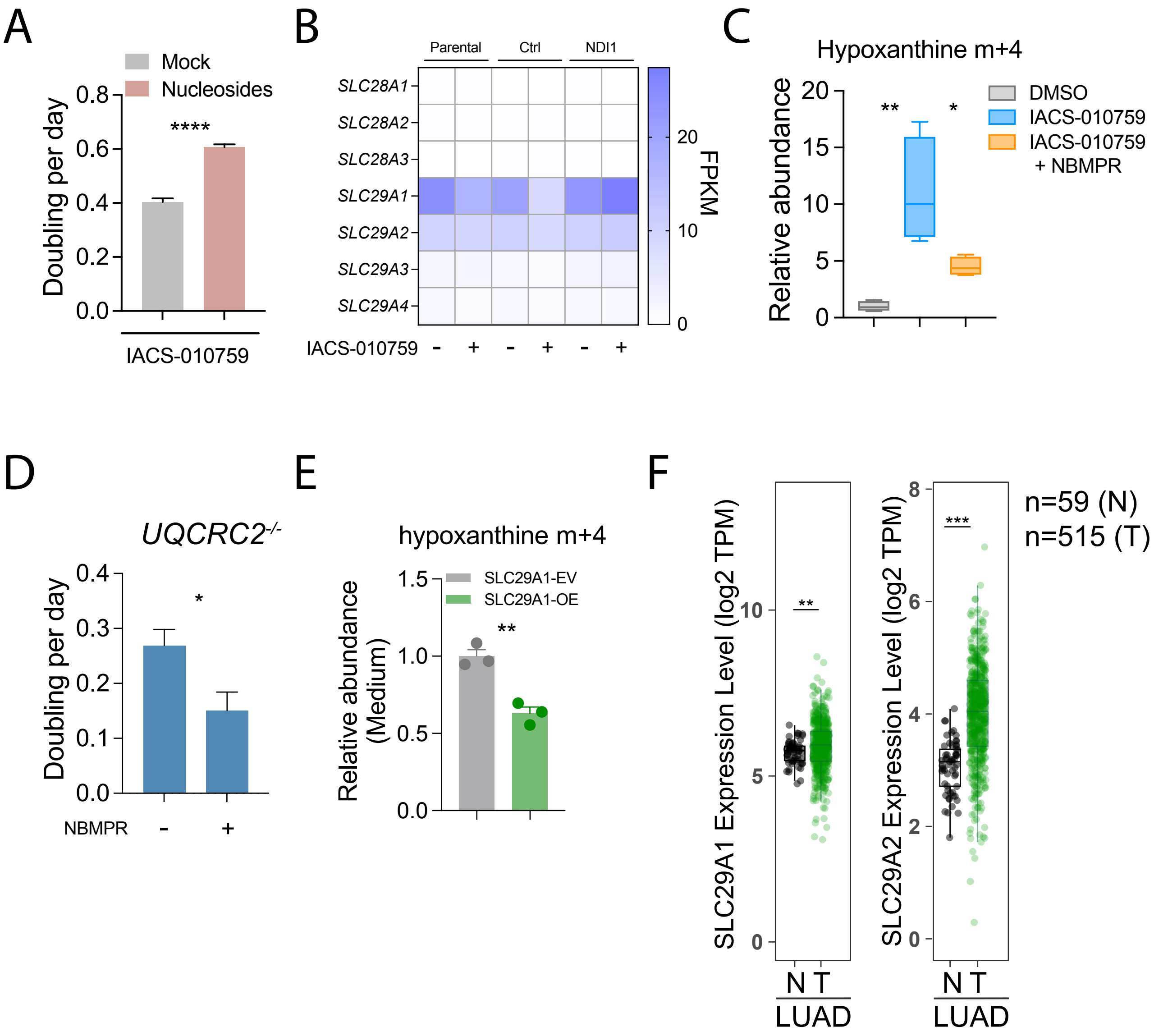
